# Spatially organized developmental cell states link paraganglioma and neuroblastoma through chromaffin-neuroblast plasticity

**DOI:** 10.64898/2026.04.07.715303

**Authors:** Jiacheng Zhu, Wenyu Li, Valeriy Paramonov, Peng Cui, Valentin Poltorachenko, Jorrit Stada, Maria Arceo, Petra Bullova, Monika Plescher, Karolina Solhuslokk Hose, Per Kogner, Jakob Stenman, Catharina Larsson, Igor Adameyko, Michael Mints, Oscar C. Bedoya-Reina, C. Christofer Juhlin, Arthur S. Tischler, Susanne Schlisio

## Abstract

Neuroblastoma (NB) and paraganglioma (PPGL) arise from the sympathoadrenal lineage, yet their developmental relationship and heterogeneity remain unclear. Single-cell and spatial transcriptomics revealed shared, spatially organized developmental states, including populations characteristic of the other tumor type, chromaffin-like cells in NB and neuroblast-like cells in PPGL and hybrids co-expressing adjacent states. Deconvolution of an independent PPGL cohort associated metastatic disease most strongly with chromaffin hybrid states, alongside connecting progenitor-like, cycling neuroblast, and early chromaffin identities, whereas non-metastatic tumors were enriched for differentiated late chromaffin identities. In mice, combined loss of the candidate 1p36 tumor suppressor *KIF1Bβ* and *NF1* recapitulates these developmental architectures by prolonging developmental plasticity, reactivating embryonic neurogenic programs, and driving chromaffin-to-neuroblast transitions generating pheochromocytoma, neuroblastoma and composite tumors. Mouse tumors contained discrete spatial domains of developmental and neoplastic states that mirrored those in human PPGL. Together, our findings establish chromaffin-neuroblast plasticity as a mechanism of sympathoadrenal tumor heterogeneity and identify hybrid developmental states as a potential indicator of plasticity associated with metastatic PPGL.

---

Neuroblastoma (NB) and pheochromocytoma/paraganglioma (PPGL) are developmentally related sympathoadrenal tumors with markedly different clinical characteristics. Neuroblastoma predominantly affects children and displays striking clinical heterogeneity, ranging from spontaneous regression to metastatic, therapy-resistant disease. In contrast, PPGLs occur predominantly in adults and are usually non-metastatic at diagnosis, although their risk of metastatic progression remains difficult to predict^1,2^. Despite their distinct clinical profiles, both neuroblastoma and PPGL derive from the same embryonic origin: the sympathoadrenal lineage, which in turn originates from transient neural crest cells. Rather than arising through a strictly linear differentiation hierarchy ^3,4^, chromaffin cells and intra-adrenal sympathoblasts (neuroblasts) emerge from multipotent Schwann cell precursors (SCPs) that spread with peripheral preganglionic motor and sensory nerves to the adrenal anlage ^5^. These SCPs, multipotent descendants of neural crest cells, give rise to chromaffin cells independently of early sympathetic ganglia development ^5,6^. More recently, single-cell transcriptomic studies identified a population of intra-adrenal immature sympathoblasts that are capable of bidirectional transitions with local chromaffin cells^7^. These bidirectional fate transitions between chromaffin cells and sympathoblasts suggest a high degree of plasticity within the sympathoadrenal lineage during development.

These observations raise the possibility that developmental cell-state transitions contribute to tumor heterogeneity. Consistent with this concept, recent transcriptomic studies have revealed substantial cellular heterogeneity in neuroblastoma ^7–12^, while PPGLs exhibit substantial transcriptional diversity associated with different genetic backgrounds and clinical outcomes ^13^. Rare composite tumors containing both PPGL and neuroblastoma components further suggest a close developmental relationship between these malignancies^14^. However, whether chromaffin-neuroblast plasticity directly contributes to tumor initiation, progression, and intratumoral heterogeneity remains unknown.

To investigate the mechanisms underlying sympathoadrenal tumor heterogeneity, we combined single-cell and spatial transcriptomics of human neuroblastoma and PPGL with genetic modeling of sympathoadrenal tumorigenesis in mice. We identify chromaffin-like, connecting progenitor-like, hybrid, and neuroblast-like developmental cell states that are shared between neuroblastoma and PPGL and spatially organized within both tumor types. Using an *NF1/KIF1Bβ*-deficient mouse model, we demonstrate that loss of *KIF1Bβ* cooperates with *NF1* deficiency to prolong developmental plasticity, promote chromaffin-to-neuroblast lineage transitions, and recapitulate these developmental cell states, giving rise to pheochromocytoma, neuroblastoma and composite tumors. Together, our findings establish developmental cell-state plasticity as a fundamental mechanism shaping heterogeneity across neuroblastoma, PPGL and composite tumors.

## Results

### Human neuroblastoma and PPGL share developmental cell states along the sympathoadrenal lineage

To characterize shared and distinct tumor cell states in human neuroblastoma (NB) and pheochromocytoma/paraganglioma (PPGL), we analyzed a Smart-seq2 single-cell RNA-sequencing cohort comprising 23 neuroblastomas and 9 PPGLs, including both newly generated and previously published datasets^12,15^ (Fig. 1a–c; Extended Data 1a–d; Supplementary Data 1). Unsupervised clustering identified tumor, stromal and immune cell populations based on established lineage markers (Fig. 1a–c). PPGL tumor clusters expressed canonical neuroendocrine markers including *CHGA, CHGB, RET, DLK1* and *PENK*, whereas neuroblastoma tumor populations expressed *PRPH, SOX4, SOX11, STMN4, ELAVL4, GAP43, MYCN* and *ALK* (Fig. 1c; Extended Data 1d; Supplementary Data 2). Additional clusters corresponded to endothelial, mesenchymal, macrophage, cortical, glia/sustentacular and immune cell populations (Extended Data 1d). A complementary Smart-seq2 dataset of healthy human adrenal glands was analyzed to define postnatal chromaffin cell states and served as a reference for subsequent deconvolution analyses (Extended Data 1e–i).

**Fig. 1.**
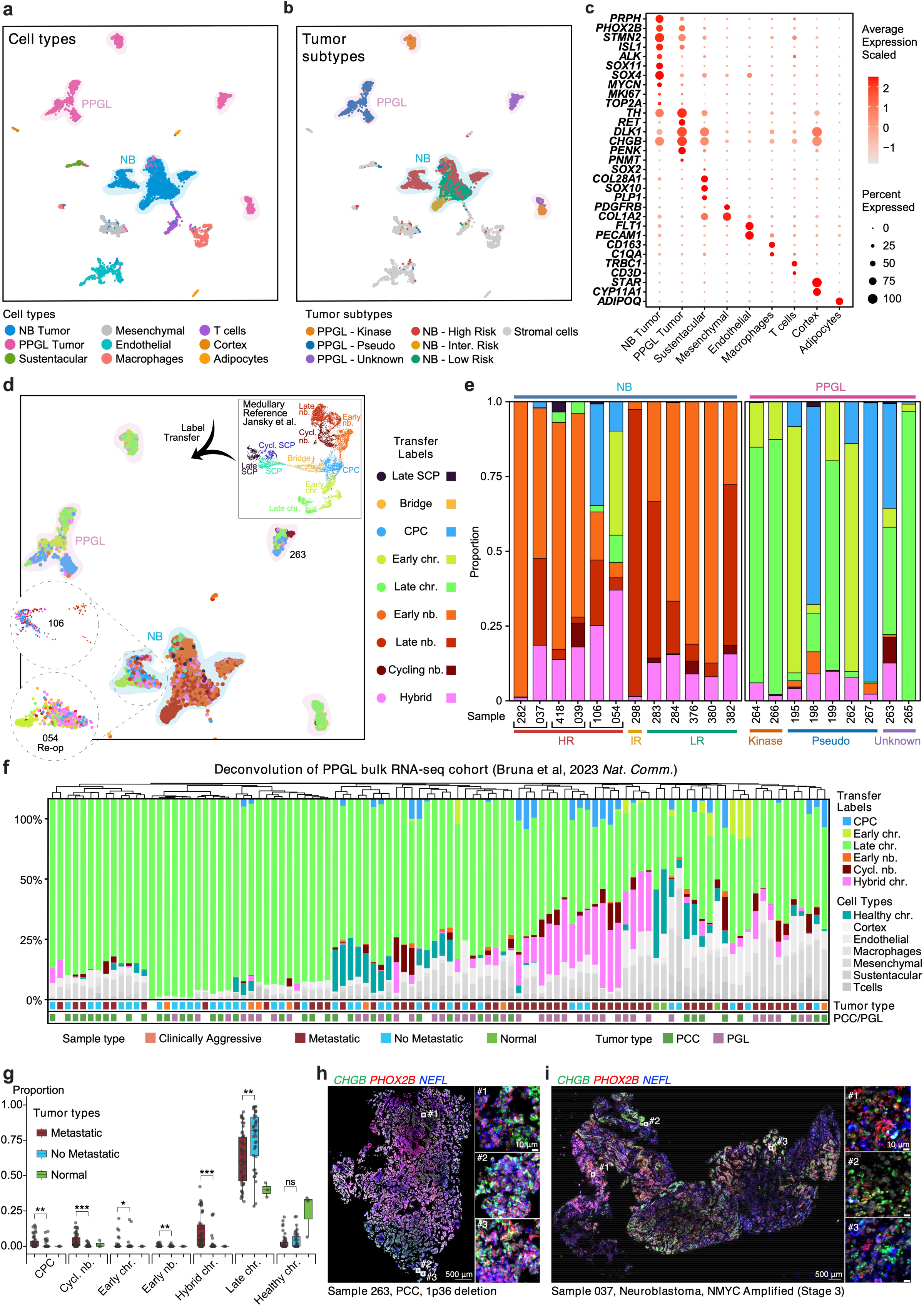
Joint single-nucleus transcriptomic analysis of human neuroblastoma and paraganglioma reveals developmental cell-state relationships. **a-c** Joint snRNA-seq analysis of human neuroblastoma (NB) and paraganglioma (PPGL). **a** UMAP representation of the merged PPGL and NB single-nucleus transcriptomic dataset colored by annotated cell type. The corresponding unsupervised clustering is shown in Extended Data 1a. **b** UMAP of the same dataset colored by tumor molecular subtype, including NB risk groups and PPGL molecular classes; non-tumor populations are grouped as stromal cells. **c** Dot plot showing canonical marker genes used for cell-type annotation. Dot color indicates scaled average expression and dot size indicates the percentage of cells expressing each gene. **d,e** Mapping of NB and PPGL tumor cells to fetal adrenal medullary developmental states by label transfer. **d** UMAP of tumor cells colored by transferred labels from the human fetal adrenal medullary reference atlas^8^, shown in the inset. Enlarged views of matched primary (sample 106) and recurrent (sample 054, obtained at reoperation) tumors highlight the change in inferred developmental cell-state composition. **e** Developmental cell-state composition of individual NB and PPGL tumor samples. Matched samples derived from the same patient are connected by black brackets. Tumor subtype and molecular classification are indicated by the annotation bars below the plot. **f,g** Deconvolution of a published PPGL bulk RNA-seq cohort^16^, comprising normal adrenal medulla, non-metastatic, clinically aggressive and metastatic PPGLs, using the developmental cell-state labels assigned to our Smart-seq2 PPGL reference dataset. **(f)** Stacked bar plot showing the inferred cell-state composition of individual PPGL samples. Sample classification (normal, non-metastatic, metastatic or clinically aggressive) and tumor type indicated below each sample. Blank annotation bars indicate unavailable sample classification or tumor type information. **(g)** Quantification of selected developmental and postnatal chromaffin/neuroblast-related cell states across normal adrenal medulla, non-metastatic PPGL and metastatic PPGL. Box plots show the median and interquartile range, with points representing individual samples. Statistical significance was assessed using two-sided Wilcoxon rank-sum tests comparing metastatic and non-metastatic PPGLs, followed by Benjamini–Hochberg false-discovery-rate (FDR) correction. ns, FDR ≥ 0.05; *FDR < 0.05; **FDR < 0.01; ***FDR < 0.001. **h,i** RNAscope validation of *CHGB* (green), *PHOX2B* (red) and *NEFL* (blue) expression in a PPGL with chromosome 1p36 deletion (sample 263) **(h)** and a MYCN-amplified stage 3 neuroblastoma (sample 037) **(i)**. Scale bars, 500 μm (whole-section images) and 10 μm (insets).

We next mapped tumor cells onto a published human fetal adrenal single-cell reference atlas^8^ using transcriptional nearest-neighbor label transfer to assign developmental identities (Fig. 1d–e). NB tumor cells were predominantly assigned to embryonic early neuroblast and late neuroblast identities, whereas PPGL cells mapped primarily to connecting progenitor-like (CPC), early chromaffin and late chromaffin states. Interestingly, both tumor types also contained hybrid developmental cell states (Fig. 1d–e), defined as tumor cells exhibiting substantial transcriptional similarity to multiple developmental reference populations rather than a single developmental identity (Extended Data 2a–e). These hybrid states likely represent cells with mixed developmental programs, consistent with incomplete stabilization of developmental identity. Moreover, NB and PPGL contained smaller populations corresponding to developmental identities of the opposite tumor type, including neuroblast-like cells within PPGL and chromaffin-like cells within NB (Fig. 1d-e). Notably, a longitudinal MYCN-amplified NB case (#106) shifted from a tumor enriched in neuroblast and chromaffin-neuroblast hybrid states at diagnosis to one enriched in chromaffin-like identities (#054) with predominantly chromaffin-associated hybrid states at reoperation (Fig. 1d-e; Extended Data 2d-e).

To determine whether developmental cell-state composition was associated with clinical behavior, we deconvolved an independent published cohort of human PPGL bulk RNA data set^16^ using the developmental labels transferred from the fetal adrenal reference atlas (Fig. 1f-g; Extended Data 2f). Metastatic PPGLs contained increased proportions of embryonic connecting progenitor (CPC)-like, cycling neuroblast, and early chromaffin-like hybrid cell states, whereas non-metastatic tumors exhibited significantly higher proportions of embryonic late chromaffin-like identities (Fig. 1f–g). Although late chromaffin-like cells represented the predominant developmental state in both metastatic and non-metastatic PPGLs, they were significantly more abundant in non-metastatic tumors. In contrast, postnatal healthy chromaffin signatures were largely restricted to normal adrenal tissue, consistent with terminal chromaffin differentiation (Fig. 1f–g). These findings indicate that metastatic PPGLs retain immature developmental cell states that are largely absent from non-metastatic tumors, whereas non-metastatic tumors remain predominantly composed of late chromaffin-like cells.

Multiplex RNAscope further validated the presence of neuroblast-like regions in PPGL and chromaffin-like regions in neuroblastoma. PHOX2B-positive regions in PPGL showed reduced CHGB and increased NEFL expression, whereas neuroblastomas contained PHOX2B-positive regions with high CHGB and low NEFL. Cells co-expressing CHGB and NEFL were also detected (Fig. 1h–i; Extended Data Fig. 2g–h).

Together, these analyses demonstrate that human neuroblastoma and PPGL share conserved developmental cell states despite representing distinct tumor entities. While multiplex RNAscope validated reciprocal chromaffin-like and neuroblast-like tumor populations using a limited set of lineage markers, it could not comprehensively resolve the full spectrum of developmental cell states or their spatial organization. We therefore next applied high-resolution spatial transcriptomics to map developmental cell states and their spatial organization in PPGL.

### Spatial transcriptomics of human PPGL reveals conserved sympathoadrenal developmental states underlying tumor heterogeneity

Having identified early and late chromaffin-like, connecting progenitor (CPC)-like, neuroblast-like, and hybrid developmental states in human NB and PPGL by single-cell transcriptomic analysis (Fig. 1), we next used Visium HD spatial transcriptomics to determine how these transcriptionally defined states are spatially organized within intact tumor tissue. We analyzed two PPGL specimens with distinct clinical and histological characteristics, including a pheochromocytoma displaying marked intratumoral heterogeneity (PPGL #1, Fig. 2a-g; Extended Data 3) and a malignant PPGL that was included in the single-cell RNA sequencing (PPGL #263 Fig. 2i-n; Extended Data 4).

**Fig. 2.**
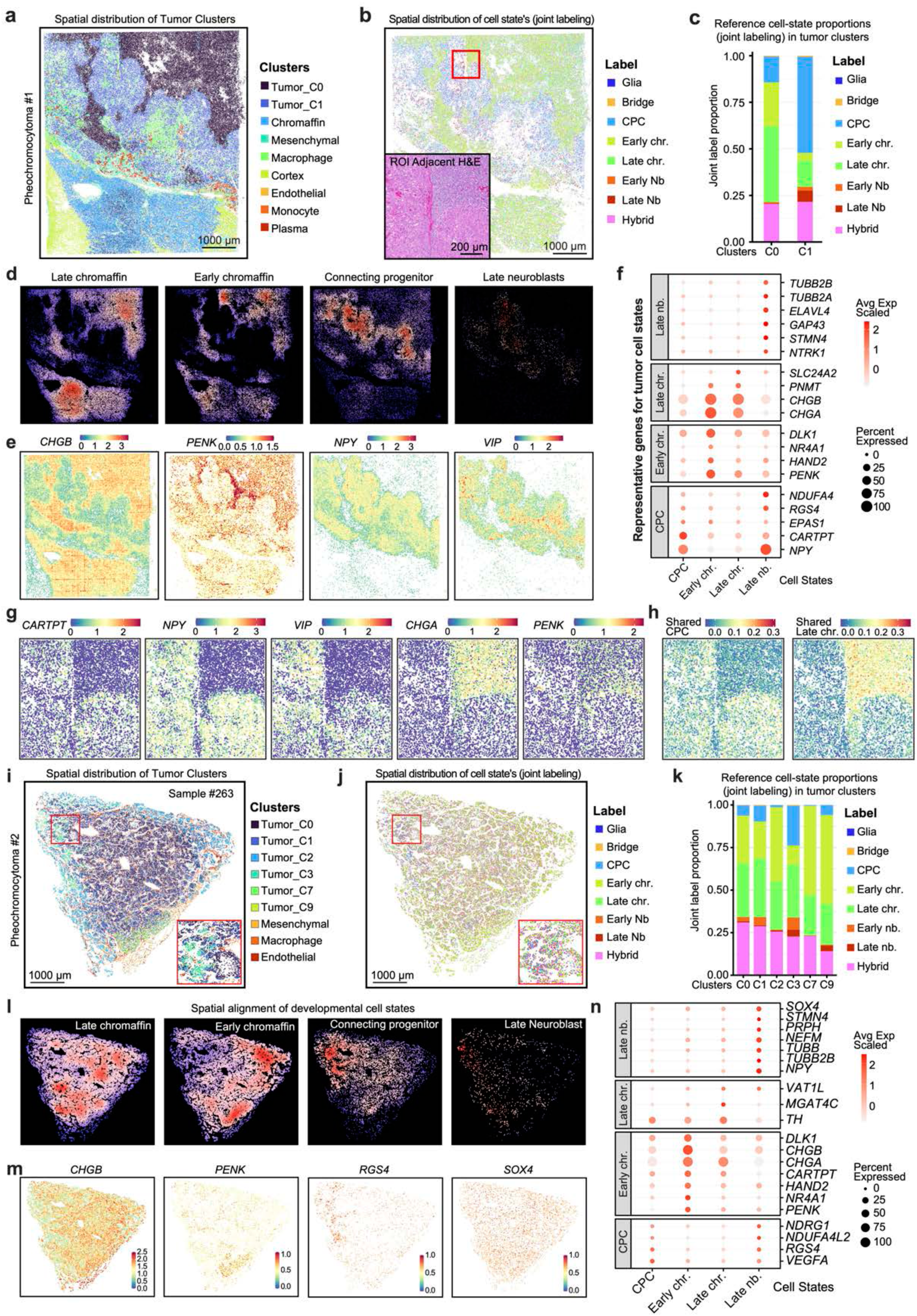
Visium HD spatial transcriptomics reveals spatial organization of developmental cell states and lineage heterogeneity in human PPGL tumors. **a-h** Visium HD spatial transcriptomic analysis of a human PPGL #1. **a** Spatial distribution of annotated tumor and non-tumor clusters across the tissue section. Clusters were annotated using canonical marker expression together with histopathological assessment. Unsupervised clustering results and marker gene expression are shown in Extended Data 4. **b** Spatial distribution of developmental cell-state assignments following label transfer from the human fetal adrenal medullary reference atlas (Jansky *et al*., 2021). Tumor cells are colored according to their assigned developmental state. The red box indicates the ROI shown in (**g-h**). The inset shows the H&E-stained adjacent tissue section corresponding to the ROI. Scale bar, 200 μm. **c** Developmental cell-state composition of Tumor_C0 and Tumor_C1, showing the proportion of cells assigned to each fetal adrenal medullary reference state. **d** Spatial density maps of reference-derived developmental cell states. Color intensity indicates relative density within each panel, from low (blue) to high (red). **e** Spatial expression of representative genes including *CHGB*, *PENK*, *NPY*, and *VIP*. **f** Dot plot summarizing expression of representative marker genes across the identified tumor cell states. Dot color indicates scaled average expression, and dot size indicates the percentage of cells expressing each gene. **g** Spatial expression of representative cell-state marker genes within the ROI (Fig. 2b, inset). **h** Spatial distribution of shared CPC and shared late chromaffin gene module scores across the ROI (Fig. 2b, inset). **i-n** Visium HD spatial transcriptomic analysis of an independent malignant PPGL (#263). **i** Spatial distribution of annotated tumor and non-tumor clusters across the tissue section. The inset highlights a region exhibiting spatially heterogeneous tumor clusters. **j** Spatial distribution of developmental cell-state assignments following label transfer from the fetal adrenal medullary reference atlas (Jansky *et al*., 2021). Tumor cells are colored according to their assigned developmental state. The inset shows a higher-magnification view of the highlighted region. **k** Developmental cell-state composition of the annotated tumor clusters. **l** Spatial density maps of reference-derived developmental cell states. Color intensity indicates relative density within each panel, from low (blue) to high (red). **m** Spatial expression of representative genes including *CHGB*, *PENK*, *RGS4*, and *SOX4*. **n** Dot plot summarizing expression of representative marker genes across the identified tumor cell states. Dot color indicates scaled average expression, and dot size indicates the percentage of cells expressing each gene.

Spatial transcriptomic analysis of PPGL #1 resolved two major tumor populations (C0 and C1) together with non-neoplastic tissue compartments (Fig. 2a; Extended Data 3a–h; Supplementary Data 3). These tumor populations corresponded to morphologically distinct regions, generating a strikingly biphasic appearance within the tumor (Fig. 2b; Extended Data 3a–b). Joint transcriptional mapping to a human fetal adrenal reference atlas revealed that tumor cluster C0 was enriched for early and late chromaffin identities, whereas tumor cluster C1 showed a larger contribution from CPC, late neuroblast and hybrid populations (Fig. 2b–c; Extended Data 3i). Hybrid-cell composition differed substantially between tumor regions (Extended Data 3g–k). Mapping developmental labels back onto the tissue section demonstrated that late and early chromaffin-like, CPC-like, and late neuroblast-like populations occupied adjacent yet spatially distinct domains rather than being intermixed (Fig. 2b,d). Representative cell-state markers and reference-derived cell-state signatures independently supported the spatial assignments (Fig. 2e–h; Extended Data Fig. 3l; Supplementary Data 4–5). Notably, PENK, an early chromaffin marker in the developmental reference, localized between CPC-like and late chromaffin-like domains, spatially mirroring the developmental progression from CPC through early to late chromaffin states (Fig. 2e,g). In contrast to PPGL #1, malignant PPGL #263 retained spatial organization of developmental cell states but exhibited greater intermixing between these states (Fig. 2i–n; Extended Data 4). Joint mapping to the fetal adrenal reference atlas identified early and late chromaffin-, CPC-, hybrid-, and neuroblast-like populations distributed throughout the tumor (Fig. 2j–n; Extended Data 4i–j, Supplementary Data 6–7). Although these developmental cell states showed greater intermixing than in PPGL #1, they remained spatially organized into neighboring domains rather than being randomly distributed throughout the tissue. The inferred developmental identities were further supported by the spatial expression of representative marker genes and their corresponding reference cell-state signatures (Fig. 2m–n; Extended Data 4).

Together, these analyses demonstrate that human PPGLs contain spatially organized developmental cell states corresponding to chromaffin-, CPC-, hybrid-, and neuroblast-like identities. Although the degree of spatial segregation varied between tumors, these developmental cell states consistently occupied discrete spatial domains within intact tumor tissue. Hybrid populations co-expressing adjacent developmental programs suggest the existence of intermediate developmental states. However, the presence of hybrid states alone does not establish developmental plasticity, which requires evidence that cells transition between developmental states over time. We therefore next investigated tumor development across embryonic, neonatal, pre-neoplastic, and established tumor stages in genetically engineered mouse models to determine whether these developmental cell states arise through lineage plasticity during tumorigenesis.

### Loss of KIF1Bβ enhances NF1-driven sympathoadrenal tumorigenesis, promoting pheochromocytoma, neuroblastoma, and composite tumors

*KIF1Bβ* is a candidate 1p36 tumor suppressor required for sympathoblast differentiation and apoptosis during sympathetic nervous system development ^17,18^. To investigate its role in sympathoadrenal tumorigenesis, we generated mice with conditional deletion of *KIF1B*β in the sympathoadrenal lineage using *DBH*-Cre^17^ and crossed them with *NF1* conditional knockout mice^19^ to generate *NF1/KIF1Bβ* double conditional knockout animals (DKO). *KIF1Bβ*cKO mice (n = 14) did not develop tumors, whereas combined loss of *KIF1Bβ* markedly potentiated NF1-driven tumorigenesis. DKO mice (n = 20) developed large adrenal tumors with complete penetrance and significantly reduced survival compared with *NF1c*KO mice (median survival 407 vs. 496 days, log-rank p = 0.0053; Fig. 3a-b; Supplementary Table 1).

**Fig. 3.**
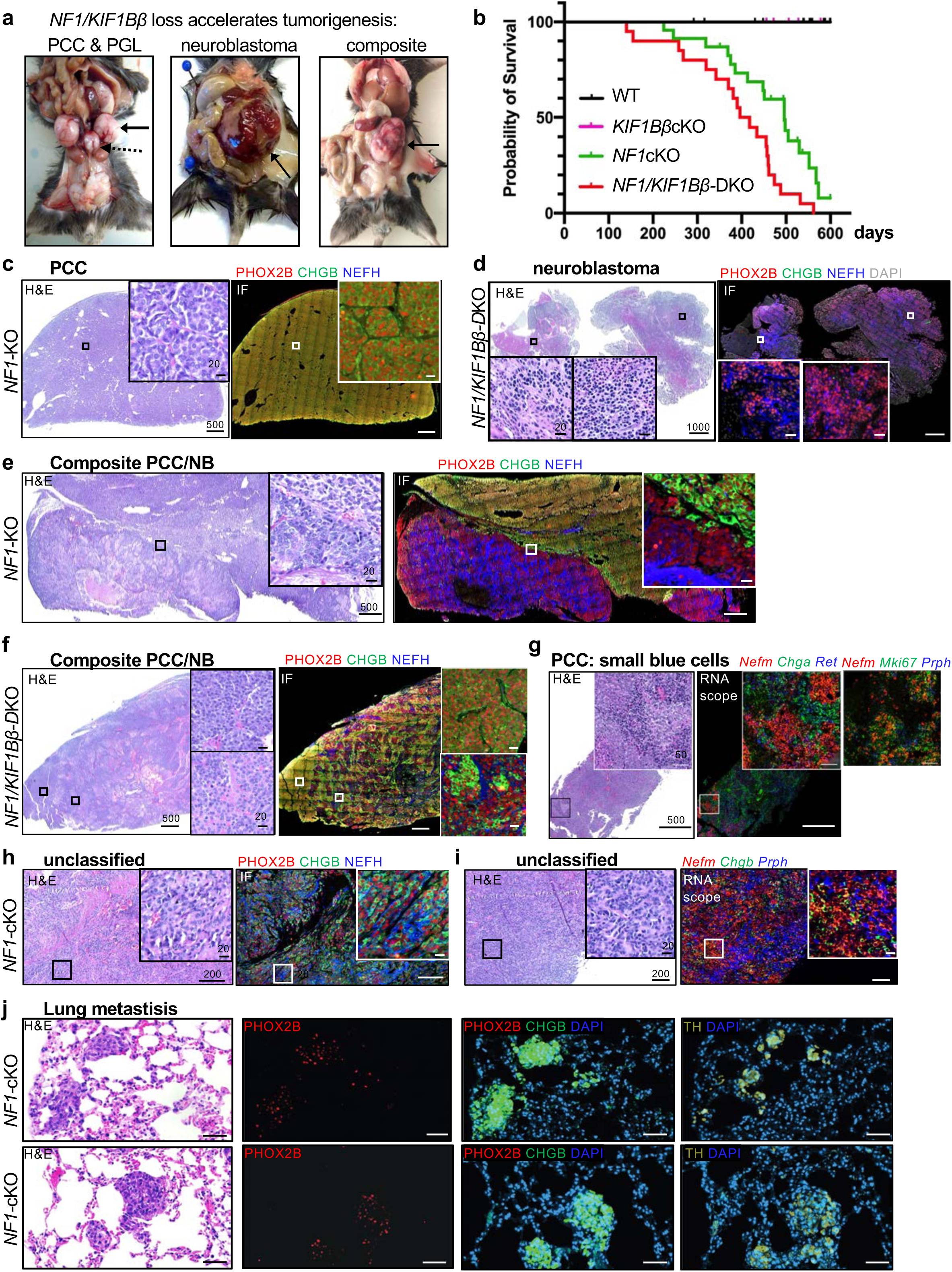
Combined *KIF1Bβ* and *NF1* loss generates pheochromocytoma, neuroblastoma, and composite tumors and accelerates tumorigenesis. **a** Gross autopsy images from tumor-bearing mice showing adrenal-derived pheochromocytoma (PCC), neuroblastoma (NB), composite PCC-NB tumors, and extra-adrenal paraganglioma (PGL) (arrows). **b** Kaplan-Meier analysis of tumor-associated survival. *NF1/KIF1Bβ*-cKO (DKO, red, n = 20) mice exhibit significantly reduced survival compared with *NF1*-cKO mice (green, n = 23), whereas *KIF1Bβ*-cKO (pink, n = 14) and WT (black, n = 37) mice remain tumor-free during the observation period. Median survival was 407 days for DKO mice and 496 days for *NF1*-cKO mice (log-rank Mantel–Cox test, P = 0.0053). **c** Representative PCC from an *NF1*-cKO mouse showing classic Zellballen-like chromaffin tumor architecture and strong PHOX2B/CHGB expression without NEFH. **d** Neuroblastoma from an DKO mouse showing small round blue cell morphology with strong PHOX2B and NEFH expression and absence of CHGB, consistent with neuroblast identity. Additional marker analyses are shown in Extended Data 1a–b. **e** Composite PCC-NB tumor from an NF1-cKO mouse with spatially segregated chromaffin-like and neuroblast-like regions. **f** Composite PCC-NB tumor from an NF1/KIF1Bβ-cKO mouse showing patchy admixture of PHOX2B⁺, CHGB⁺, and NEFH⁺ tumor populations. **g** PCC from an NF1-cKO mouse containing focal nests of “small blue cells.” RNAscope identifies neuro-proliferative-like *Nefm⁺/Mki67⁺/Prph⁺* cells embedded within Chga/Chgb-positive PCC tissue. Extended marker analyses are shown in Extended Data 1c. **h-i** Unclassified tumor from an *NF1*-cKO mouse displaying mixed chromaffin and neuroblast features. Immunofluorescence and RNAscope reveal co-expression of PHOX2B, CHGB, NEFH (h), *Nefm*, *Chgb* and *Prph* (i), consistent with a hybrid chromaffin–neuroblast-like state. **j** Lung metastases from *NF1*-cKO mice showing PHOX2B⁺CHGB⁺TH⁺ tumor cell clusters within lung parenchyma (also see Extended Data 1e). Scalebar of overview in c: 500 μm; d: 1000um; e: 200um; f and g: 500um; zoom of boxed image: 20 μm; h and i: 400 μm, zoom of boxed image: 50 μm; j: 50 μm.

Histopathological analysis revealed a spectrum of sympathoadrenal tumors, the majority displaying features of PPGL, including Zellballen-like architecture and chromaffin marker expression (Fig. 3c; Supplementary Table 2). In addition, neuroblastoma and composite tumors containing spatially distinct PPGL-like and neuroblastoma-like regions were observed (Fig. 3d-f; Extended Data 5a–b; Supplementary Table 2). Histologically detectable lung metastases were observed in a subset of *NF1*cKO and DKO animals (Fig. 3j; Extended Data 5c).

Notably, many PPGLs contained focal aggregates of small round blue cells lacking chromaffin markers but expressing neuroblast-associated markers and MKI67, indicating proliferative neuroblast-like populations embedded within otherwise chromaffin tumors (Fig. 3g; Extended Data 5d). In addition, an unclassified tumor contained cells co-expressing chromaffin and neuroblast markers, consistent with a hybrid chromaffin-neuroblast identity (Fig. 3h–i; Extended Data 5e). Together, these findings demonstrate that NF1/KIF1Bβ-deficient tumors comprise chromaffin-like, neuroblast-like, and hybrid populations, indicating substantial cellular heterogeneity within these sympathoadrenal tumors.

### Aberrant chromaffin-neuroblast lineage transitions drive neuroblast expansion in NF1 and NF1/KIF1Bβ mutant adrenals

To determine whether PPGL and neuroblastoma arise through altered developmental lineage trajectories, we performed single-cell transcriptomic profiling across embryonic, pre-neoplastic, and tumor stages. Embryonic sympathoadrenal progenitors were lineage-traced using tamoxifen-inducible *Sox10*CreERT2;Rosa26-YFP mice, with tamoxifen administered at E11.5 and adrenal glands harvested at E17.5 for Smart-seq2 analysis (Extended Data 6a). Because embryonic tamoxifen induction limited postnatal survival, later stages were analyzed using the constitutive *DBH*-Cre driver. At E17.5, unsupervised clustering resolved eleven transcriptionally distinct medullary populations, revealing remarkable heterogeneity within the embryonic sympathoadrenal lineage. These included SCPs, bridge cells, chromaffin cells, multiple neuroblast populations, and hybrid chromaffin-neuroblast intermediates (Fig. 4a-c; Supplementary Data 9). Cluster identities were supported by lineage-specific marker expression (Fig. 4b). Cluster identities were supported by established chromaffin and neuroblast markers (Fig. 4b–c). Notably, an SCP-derived bridge–neuroblast population, preferentially detected in mutant adrenals, retained bridge-associated features while expressing neuroblast markers together with *Ret, Fgf13* and *Cux2*. Transcriptional nearest-neighbor mapping to a published mouse embryonic adrenal reference^7^ confirmed cluster annotation and developmental identity across all genotypes (Fig. 4d).

**Fig. 4.**
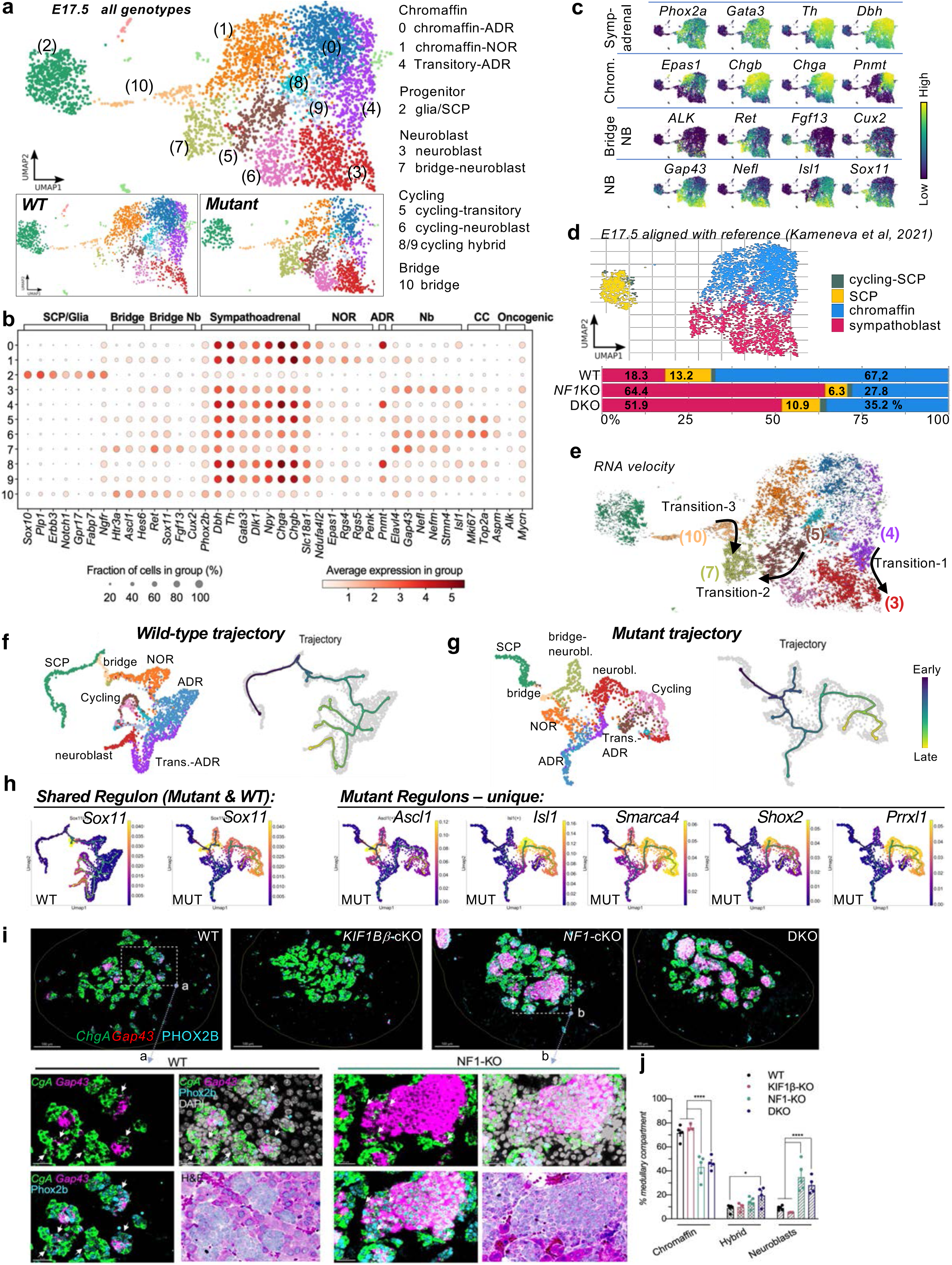
*NF1/KIF1Bb* loss induces neurogenic regulatory programs along altered sympathoadrenal developmental trajectories. **a** Cellular landscape of lineage-traced embryonic medulla. UMAP of SOX10-lineage–traced (YFP⁺) E17.5 adrenal medullary cells from WT and mutant (*NF1*-KO and DKO) embryos identifying 11 transcriptionally distinct clusters, including SCP/glial, bridge, chromaffin, neuroblast, and hybrid populations. Insets show genotype-specific distributions. **b** Lineage-associated gene expression across clusters. Dot plot showing expression of representative SCP/glial, bridge, chromaffin, neuroblast, and cell-cycle genes across clusters. **c** Feature plots of Smart-seq2 single-cell RNA sequencing from lineage-traced embryonic medulla shown. Representative lineage and state-specific markers validate cluster annotation. **d** Alignment to published sympathoadrenal reference states. Integration with a published embryonic adrenal reference dataset (Kameneva et al.) confirms annotation of SCP, chromaffin, and neuroblast states. *NF1*-KO and DKO embryos show marked enrichment of neuroblast populations compared with WT. **e** RNA velocity identifies multiple trajectories toward neuroblast identity. Velocity analysis reveals three principal neuroblast-associated transitions arising from adrenergic chromaffin, cycling transitional, and bridge cell populations, indicating enhanced neurogenic lineage bias in mutant embryos. **f**-**g** PAGA-informed trajectory analysis of WT and mutant embryos. Developmental trajectory embeddings reveal altered lineage progression and expanded neuroblast-associated trajectories in mutant embryos compared with WT. **h** Regulon activity mapped onto developmental trajectories. Shared regulons active in both WT and mutant embryos include *Sox11* (*Sox4*, *EPAS1* and *Phox2a* shown in Extended Data 2c), Mutant-specific regulons include *Ascl1*, *Isl1*, *Smarca4, Shox2* and *Prrxl1* (*Hand1*, *Hand2* and *Npdc1* shown in Extended Data 2d) are strongly enriched along neuroblast and bridge-neuroblast trajectories and absent in WT embryos. **i** *In situ* validation of neuroblast expansion in mutant embryos. Combined RNAscope (*CgA, Gap43*) with immunofluorescence (PHOX2B) on E17.5 adrenal sections reveal marked expansion of PHOX2B⁺/*Gap43⁺/CgA*⁻ neuroblast clusters in *NF1*-KO and DKO embryos compared with WT and *Kif1bβ*-KO controls. Rare hybrid cells co-expressing chromaffin and neuroblast markers (PHOX2B⁺/*Gap43⁺/CgA*⁺; arrows) are predominantly located at the periphery of neuroblast clusters. **j** Quantification of chromaffin, hybrid, and neuroblast populations across genotypes (two-way ANOVA with Tukey’s correction). Scale bars for main panels and a/b insets: 100 and 30 μm, respectively.

Comparison of wild-type (WT) and mutant embryos revealed marked redistribution of developmental cell states rather than the emergence of novel populations (Fig. 4a insets, 4d; Extended Data 6b). WT adrenals were dominated by chromaffin cells, whereas *Nf1*-KO and DKO embryos exhibited a pronounced expansion of bridge-neuroblast and neuroblast populations accompanied by a reduction in chromaffin cells (Extended Data 6b).

Hybrid populations were identified across all genotypes, consistent with transient intermediate states along the chromaffin-neuroblast developmental continuum. Bridge-neuroblast population (cluster 7) expressed bridge and neuroblast markers with low *Th* expression, consistent with a transitional neurogenic state (Fig. 4b–c; Supplementary Data 9).

To assess the directionality of lineage relationships underlying these state shifts, we performed RNA velocity analysis on the E17.5 dataset (Fig. 4e). Velocity vectors were preferentially directed from bridge and chromaffin-associated intermediates toward neuroblast populations, indicating increased progression along neuroblast-directed developmental trajectories in *Nf1*-KO and DKO embryos.

Taken together, these findings demonstrate that *Nf1* deficiency and *Nf1/Kif1bβ* deficiency does not generate novel sympathoadrenal cell types but instead redistributes cells toward neuroblast-directed developmental trajectories. This shift results in expansion of proliferative and bridge-associated neuroblast populations at the expense of chromaffin differentiation, revealing enhanced developmental plasticity within the embryonic adrenal medulla.

### *Nf1* loss, either alone or in combination with *Kif1bβ* loss, induces neurogenic regulatory programs

To define the transcriptional mechanisms underlying the altered developmental trajectories, we performed trajectory inference and SCENIC regulon analysis on the E17.5 single-cell datasets (Fig. 4f-h; Extended Data 6c-g). Trajectory analysis recapitulated the canonical sympathoadrenal differentiation continuum in wild-type embryos, progressing from SCPs through bridge intermediates toward chromaffin cells with a minor branch extending toward neuroblast states (Fig. 4f). In contrast, *Nf1*-cKO and DKO embryos displayed expansion of neuroblast-associated trajectories together with strengthened chromaffin-to-neuroblast and bridge-to-neuroblast branches, consistent with the RNA velocity analysis (Fig. 4g).

Mapping regulon activity onto developmental trajectories showed that *EPAS1* activity was confined to differentiated noradrenergic chromaffin cells, whereas *SOX4* and *SOX11* increased along trajectories toward bridge-neuroblast and neuroblast populations (Fig. 4h; Extended Data 6c-f). In contrast, *NF1*-cKO and DKO embryos selectively activated neurogenic regulons including *Ascl1, Isl1, Hand1, Hand2, Prrxl1, Shox2*, and *Smarca4*, whereas the neuronal repressor *Rest* remained preferentially active in wild-type chromaffin populations (Fig. 4h; Extended Data 6d-f). Multiplexed RNAscope and immunofluorescence independently validated chromaffin, neuroblast and hybrid populations and confirmed marked neuroblast expansion in *Nf1*-cKO and DKO embryos (34.9% and 28.2%, respectively) compared with WT (9.1%), accompanied by a reciprocal reduction in chromaffin cells (Fig. 4i–j; Extended Data 6h).

Together, these findings demonstrate that *Nf1/Kif1bβ* loss biases embryonic sympathoadrenal trajectories toward neuroblast-like states.

### Neonatal persistence of neuroblast populations and hybrid states in NF1-deficient adrenal medulla

To determine whether the embryonic lineage imbalance persisted after birth, we examined adrenal glands from postnatal day 5 (P5) mice (Fig. 5a; Extended Data 7). Whereas wild-type adrenals consisted predominantly of chromaffin cells, *Nf1*-KO adrenals contained prominent neuroblastic islands and rare hybrid cells within the medulla (Fig. 5a; Extended Data 7a–b). Neuroblasts (TH⁻ and TH-low combined) comprised ∼15% of the medullary compartment in *Nf1*-cKO adrenals compared with 0–1% in wild-type controls (p < 0.001), accompanied by a reciprocal reduction in chromaffin cells (∼65% versus ∼90%, p < 0.0001) and persistent medullary enlargement (Fig. 5a; Extended Data 7c–d). Hybrid cells were detected in *Nf1*-cKO but not wild-type adrenals.

**Fig. 5.**
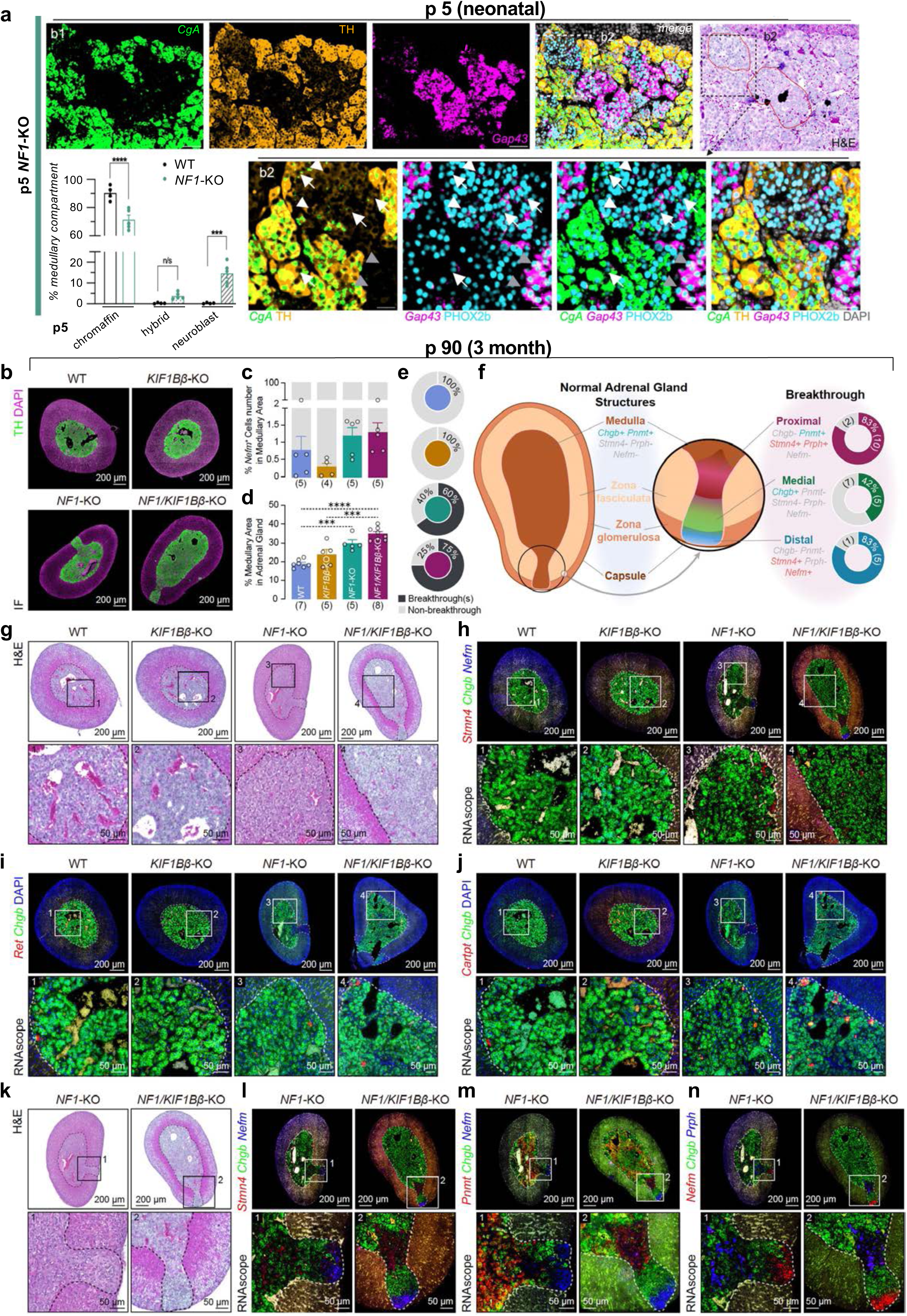
*Nf1* deficiency drives sympathoadrenal lineage plasticity and focal chromaffin-to-neuroblast lineage reversal. **a** *NF1* deficiency prolongs neuroblast-rich medullary states into neonatal life. Combined RNAscope/immunofluorescence analysis of postnatal day 5 (P5) *Nf1*-KO adrenal glands reveals persistent *Gap43*⁺ neuroblastic islands embedded within the medullary compartment compared with WT littermates (WT shown in Extended Data 3a). Overview images correspond to Extended Data 3b (b1), whereas lower panels (b2) show higher magnification views. TH-negative and TH-low neuroblast populations (PHOX2B⁺/*Gap43⁺/Chga⁻*) together with rare hybrid cells co-expressing chromaffin and neuroblast markers (PHOX2B⁺/*Gap43⁺/Chga⁺/*TH⁺; arrows) are detected within mutant adrenal glands. Quantification demonstrates increased neuroblast populations and reduced chromaffin cells in *Nf1*-KO adrenal glands compared with WT controls (two-way ANOVA with Sidak’s multiple-comparison correction). **b** Adult (P90) *Nf1/Kif1bβ* mutant adrenal glands exhibit focal cortical breakthrough lesions associated with chromaffin-to-neuroblast lineage reversal. TH immunofluorescence reveals medullary hyperplasia and focal cortical “breakthrough” lesions in *Nf1-*KO and DKO adrenal glands, in which TH⁺ medullary cells extend through the adrenal cortex toward the capsule. **c** Quantification of *Nefm*⁺ neuroblast within the adult medullary core. Neuroblasts are infrequent across genotypes, indicating that the prominent embryonic and neonatal neuroblast populations largely resolve by adulthood within the medulla. **d** Quantification of medullary area (TH⁺) relative to total adrenal gland area. *Nf1*-KO and DKO adrenal glands show significant medullary enlargement compared with WT and *Kif1bβ*-KO controls (one-way ANOVA with Tukey’s multiple-comparison test: *** *p* < 0.001 or **** *p* < 0.0001). *NF1*-KO (29.8 ± 1.8%, n = 5) and *NF1*/*Kif1bβ*-DKO mice (34.8 ± 1.2%, n = 8) compared with WT (18.9 ± 0.8%, n = 7) and *Kif1bβ*-KO controls (23.7 ± 2.8%, n = 5). **e** Frequency of cortical breakthrough lesions across genotypes based on TH⁺ area quantification. **f** Schematic model illustrating the spatial organization of cortical breakthrough lesions. **g-j** Histological and RNAscope characterization of medullary hyperplasia. The expanded medullary compartment broadly retains chromaffin marker expression (*Chgb*), while neuroblast-associated genes (*Stmn4, Nefm, Ret, Cartpt*) are detected only in scattered cells within the medullary core. **k-n** Analyses of cortical breakthrough lesions reveal spatial transitions from chromaffin marker expression (*Chgb, Pnmt*) toward neuroblast-associated programs (*Stmn4, Nefm, Prph*), consistent with focal chromaffin-to-neuroblast lineage reversal. Scale bars: b, g–n overview panels, 200 μm; inset panels, 50 μm; a main panels and inset panels, 100 μm and 30 μm, respectively.

### Postnatal medullary hyperplasia with cortical breakthroughs reveals chromaffin-to-neuroblast lineage reversal in Nf1/Kif1bβ-deficient adrenals

By 3 months, adrenal medullae across all genotypes consisted predominantly of chromaffin cells with only rare neuroblast-associated cells, indicating that the neuroblast and hybrid populations observed at P5 were transient and subsequently acquired a chromaffin fate (Fig. 5b–c,g–j; Extended Data 8a–b).

Despite restoration of chromaffin identity, mutant adrenals retained significant medullary expansion in *Nf1*-KO (29.8 ± 1.8%) and DKO (34.8 ± 1.2%) mice compared with WT (18.9 ± 0.8%) and *Kif1bβ*-KO controls (23.7 ± 2.8%; p < 0.001; Fig. 5d). Notably, most mutant adrenals (60–75%, Fig. 5e–f; Extended Data 8c) displayed focal cortical “breakthroughs” defined by TH+ medullary cells breaching the cortical boundary and extending toward the adrenal capsule. Breakthrough regions displayed spatially ordered gradients of lineage markers, progressing from chromaffin-like cells proximally to neuroblast-like cells (*Stmn4*⁺/*Prph*⁺) distally, with an intermediate *Chgb*-positive zone separating the two identities (Fig. 5i–n). RNAscope further revealed increased expression of the neurogenic receptor tyrosine kinase *Ret* within cortical breakthrough regions relative to the surrounding medullary core in all examined *Nf1*-KO (3/3) and DKO (4/4) lesions (Extended Data 8c). These transitional patterns were observed in *Nf1*-KO and DKO mice but were absent from WT and *Kif1bβ*-KO controls (Fig. 5g–h,k–n).

Together, these findings indicate that despite restoration of chromaffin identity in the adult adrenal medulla, *Nf1/Kif1bβ*-deficient cells retain latent developmental plasticity that re-emerges locally within cortical breakthrough lesions as chromaffin-to-neuroblast lineage reversal.

### Single-cell transcriptomic analysis identifies heterogeneous neoplastic chromaffin, hybrid, and neuroblast-like tumor states

To determine how developmental lineage plasticity contributes to tumor heterogeneity, we performed Smart-seq2 single-cell RNA sequencing of 3-month-old pre-neoplastic and aged tumor-bearing *Nf1/Kif1bβ*-DKO adrenals together with WT and *Kif1bβ*-KO controls (Fig. 6; Supplementary Data 8). Unsupervised clustering of 6,274 cells identified 33 populations spanning adrenal and stromal compartments (Fig. 6a).

**Fig. 6.**
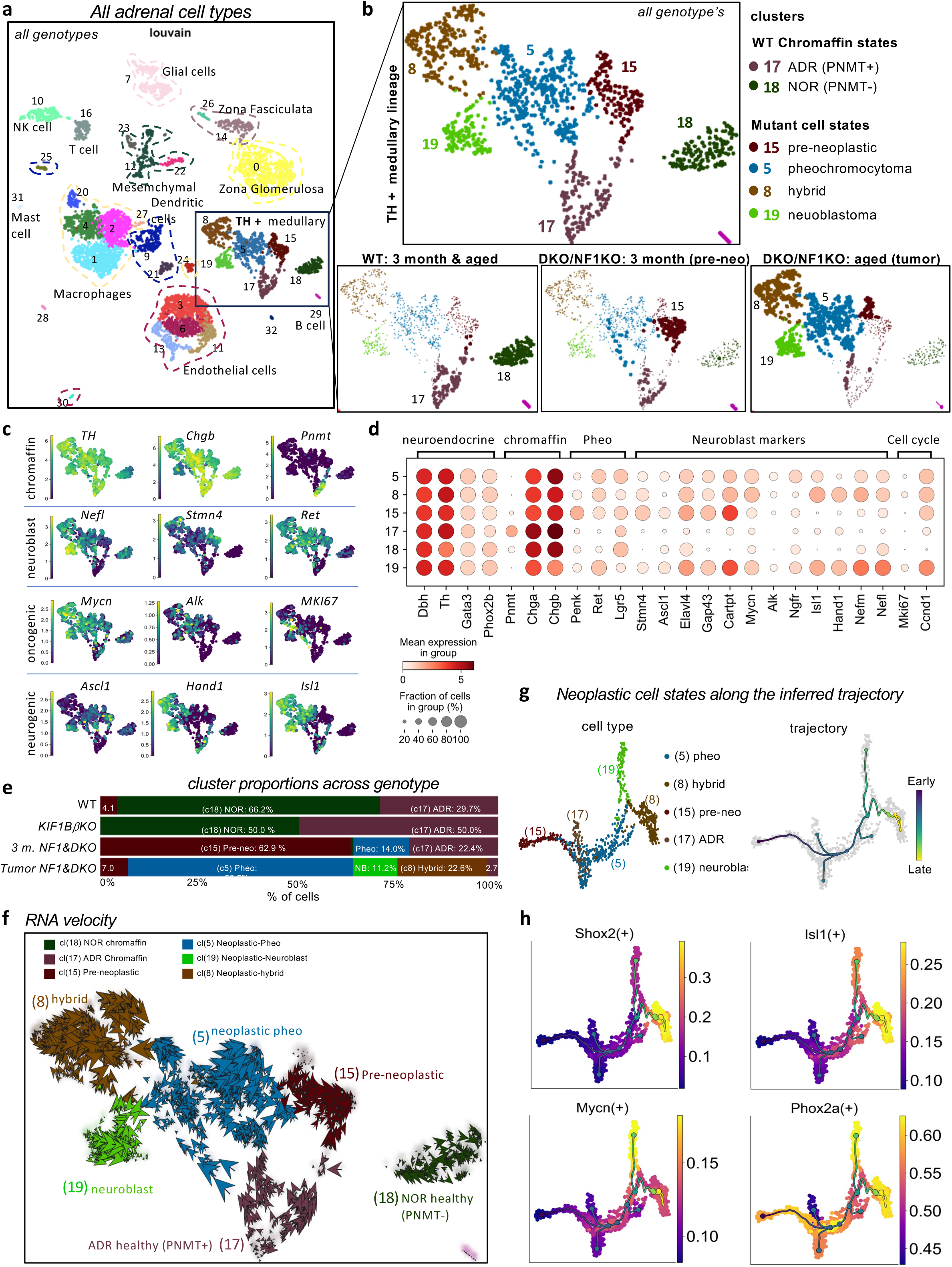
Single-cell transcriptomics reveals heterogeneous chromaffin, hybrid, and neuroblast-like tumor states in Nf1/Kif1bβ mutant adrenal glands. **a** Single-cell atlas of adrenal gland cell populations across genotypes and disease stages. UMAP embedding of 6,274 Smart-seq2–profiled adrenal cells from WT, *Kif1bβ*-KO, 3-month-old pre-neoplastic DKO, and aged tumor-bearing DKO mice, identifying medullary, cortical, endothelial, immune, and stromal compartments. **b** Identification of medullary tumor cell states and their distribution across genotypes and disease stages. TH⁺ medullary cells from panel a are shown, highlighting adrenergic chromaffin (ADR; cluster 17), noradrenergic chromaffin (NOR; cluster 18), a pre-neoplastic chromaffin population (cluster 15), and three tumor-associated states: pheochromocytoma-like (cluster 5), hybrid chromaffin–neuroblast (cluster 8), and neuroblast-like cells (cluster 19). Lower panels show genotype- and stage-specific distributions of these populations. **c** Feature plots of representative chromaffin markers (*Th, Chgb, Pnmt*), neuroblast-associated genes (*Nefl, Stmn4, Ret*), oncogenic drivers (*Mycn, Alk, Mki67*), and neurogenic transcription factors (*Ascl1, Hand1, Isl1*). **d** Dot plot summarizing lineage-associated and tumor-associated gene expression across medullary cell states. **e** Genotype- and stage-dependent shifts in medullary cell composition. WT and Kif1bβ-KO adrenal glands are dominated by chromaffin populations, whereas 3-month-old Nf1/Kif1bβ-DKO glands are enriched for pre-neoplastic cells and aged tumors display expansion of pheochromocytoma-like, hybrid, and neuroblast-like populations. **f** RNA velocity analysis suggests directional transitions from pre-neoplastic chromaffin cells toward hybrid and neuroblast-like tumor states. **g** Graph-based trajectory reconstruction resolving branching lineage relationships linking chromaffin, hybrid, and neuroblast-like tumor populations. Cells are colored by cluster identity (left) and inferred latent progression (right). **h** SCENIC regulon activity mapped onto tumor trajectories reveals activation of neurogenic regulons (*Shox2, Isl1*) in hybrid states and *Mycn*-centered regulon activity in neuroblast-like tumor cells. The core sympathoadrenal lineage regulon *Phox2a* remains broadly active across medullary tumor states despite acquisition of neurogenic programs. Additional regulons are shown in Extended Data 5f.

Analysis of the TH⁺ sympathoadrenal compartment identified adrenergic and noradrenergic chromaffin populations in WT and *Kif1bβ*-KO adrenals, whereas *Nf1*-cKO and DKO adrenals showed age-dependent expansion of pre-neoplastic and neoplastic states, including pheochromocytoma-like, hybrid chromaffin–neuroblast and neuroblastoma-like populations (Fig. 6b–e; Extended Data 9a). Compared with normal chromaffin cells, the pre-neoplastic population showed reduced chromaffin differentiation and activation of developmental and neurogenic genes, including *Gap43, Elavl4, Stmn4, Ret* and *Ascl1* (Fig. 6c–d; Supplementary Data 10). Across the neoplastic states, chromaffin differentiation progressively declined as neurogenic programs increased, with pheochromocytoma-like cells retaining chromaffin features and *Ret* expression, whereas hybrid and neuroblastoma-like states increasingly expressed *Hand1, Isl1, Shox2, Nefl* and *Mycn*, accompanied by reduced *Chgb* expression in the neuroblastoma-like state (Fig. 6c–d; Supplementary Data 10).

RNA velocity, latent time and graph-based trajectory inference revealed dynamic relationships among the neoplastic states (Fig. 6f–g; Extended Data 9b–d). Velocity vectors indicated transitions from pheochromocytoma-like toward more neuroblast-associated states, while latent time and trajectory analyses positioned these populations along a continuum of progressive neurogenic reprogramming.

SCENIC analysis revealed activation of neurogenic regulons, including *Shox2* and *Isl1*, along the tumor trajectory, with highest activity in hybrid cells (Fig. 6h; Extended Data 9e–f). *Mycn* regulon activity was selectively enriched in neuroblastoma-like cells despite broader *Mycn* expression, whereas *Phox2a* activity remained preserved across tumor states, indicating maintenance of sympathoadrenal lineage identity during neurogenic reprogramming.

Together, these analyses indicate that *Nf1/Kif1bβ*-deficient tumors undergo progressive neurogenic reprogramming, re-engaging regulatory programs active during embryonic sympathoadrenal development, with *Mycn* program activation reinforcing the neuroblastoma-like state.

### Spatial *in situ* sequencing reveals organized tumor cell states and composite architecture in *NF1/KIF1Bβ*-deficient adrenal tumors

To spatially map the developmental and neoplastic cell states identified by single-cell analysis, we performed targeted *in situ* sequencing (ISS) on 14 adrenal sections from adult and aged WT, *Kif1bβ*-KO, *Nf1*-KO and *Nf1/Kif1bβ*-DKO mice using a 100-gene panel representing embryonic, postnatal, pre-neoplastic and tumor states (Fig. 7a; Supplementary Data 11). Reference-based deconvolution reconstructed the canonical capsule–cortex–medulla organization of WT adrenals, validating the spatial mapping approach (Extended Data 10a–b; Supplementary Data 12).

**Fig. 7.**
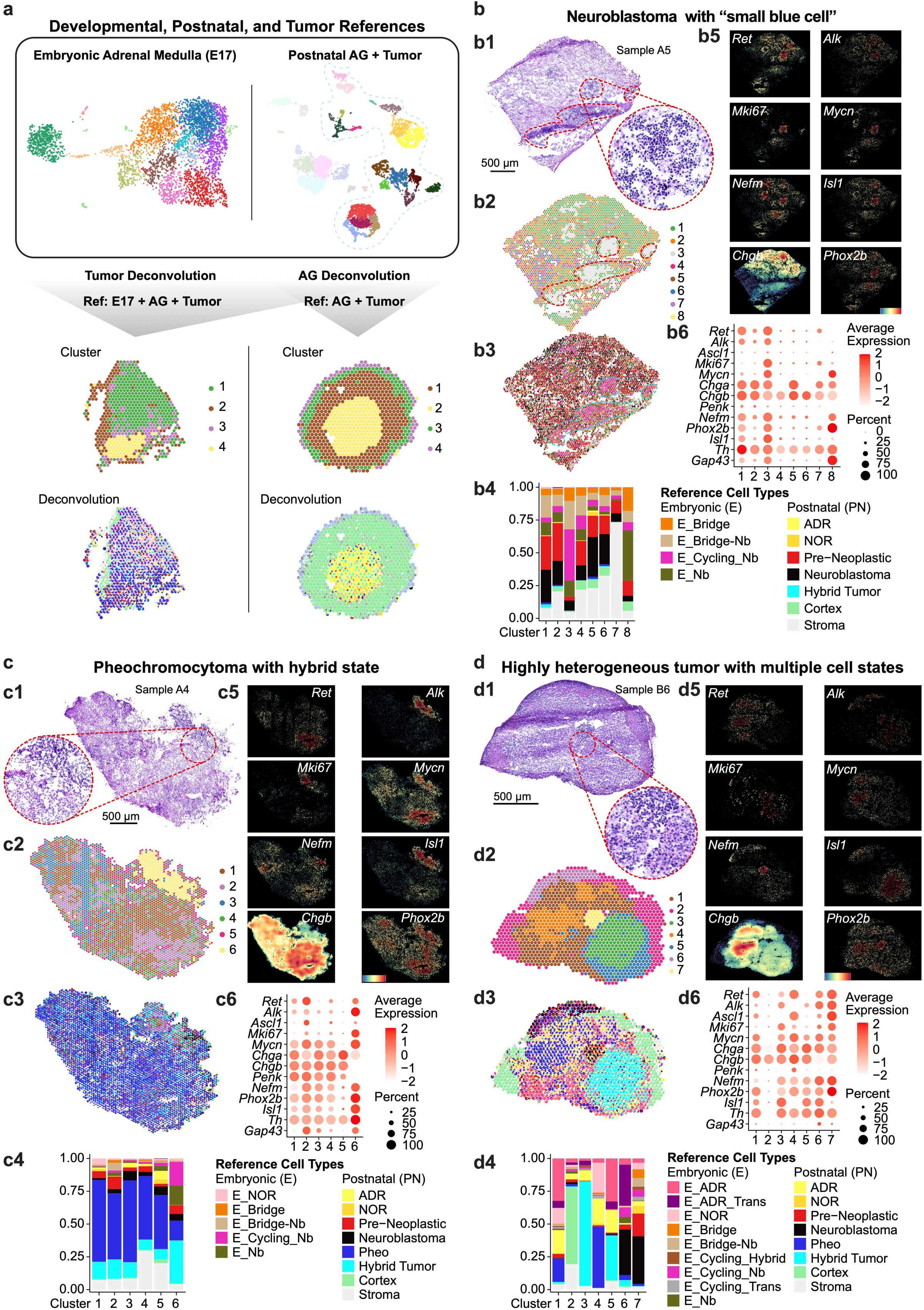
Spatial transcriptomics (ISS) reveals tumor architecture and reactivation of embryonic sympathoadrenal programs. **a** Schematic overview of the spatial deconvolution strategy. Developmental (E17 embryonic adrenal medulla), postnatal adrenal gland, pre-neoplastic, and tumor single-cell RNA-seq datasets generated in this study were used as reference atlases for spatial deconvolution of ISS profiles. **b-d** Spatial mapping of tumor cell states by targeted in situ sequencing (ISS). ISS was performed on adrenal tumors from Nf1/Kif1bβ-DKO (b–c) and Nf1-KO (d) mice using a targeted gene panel derived from embryonic, postnatal, pre-neoplastic, and tumor single-cell datasets. H&E staining illustrates tumor architecture, followed by spatial clustering, deconvolution-based cell-state mapping, marker gene expression, and cluster-level gene expression summaries. Reference states include embryonic chromaffin, embryonic neuroblast, embryonic cycling neuroblast, pre-neoplastic chromaffin, pheochromocytoma-like, neuroblastoma-like, and hybrid tumor populations. **b** Neuroblast-rich tumor region (sample A5, DKO). Spatial deconvolution reveals extensive neuroblastoma-like and embryonic neuroblast programs across the tumor. A discrete small blue cell–rich region (cluster 3) is specifically enriched for embryonic cycling neuroblast signatures together with elevated *Ret, Alk, Mycn, Isl1*, and *Mki67* expression, whereas surrounding regions show enrichment for non-cycling embryonic neuroblast, bridge-neuroblast, pre-neoplastic, and neuroblastoma-like states. **c** Pheochromocytoma with focal hybrid region (sample A4, DKO). The tumor is predominantly composed of pheochromocytoma-like states with a discrete hybrid region (cluster 6) enriched for hybrid tumor, embryonic cycling neuroblast, and pheochromocytoma-like signatures, together with elevated *Alk, Mycn, Nefm, Isl1*, and *Mki67* expression and residual chromaffin marker expression. **d** Highly heterogeneous tumor with multiple spatially distinct cell states (sample B6, *Nf1*-KO). Spatial mapping reveals coexisting pheochromocytoma-like (cluster 4), hybrid (cluster 3), and neuroblastoma-like (cluster 7) tumor regions organized into discrete spatial domains. Additional regions contain embryonic chromaffin and transitional states together with pre-neoplastic signatures, highlighting the coexistence of embryonic, pre-neoplastic and neoplastic developmental states within an established tumor.

Analysis of seven adrenal tumors revealed marked inter- and intratumoral heterogeneity, ranging from tumors dominated by pheochromocytoma-like or hybrid states to highly heterogeneous lesions containing spatially distinct pheochromocytoma-, hybrid-, neuroblastoma-like and pre-neoplastic populations (Fig. 7b–d; Extended Data 10c–f). Notably, focal regions corresponding to embryonic chromaffin and neuroblast states were also detected within established tumors, indicating reactivation of developmental programs. Hybrid regions co-expressed chromaffin and neuroblast lineage programs, providing spatial validation of the transitional states identified by single-cell analysis (Fig. 7b6,c6,d6). These developmental and neoplastic states occupied spatially organized domains within individual tumors, recapitulating the spatial organization of chromaffin-, hybrid- and neuroblast-like states observed in human PPGL (Fig. 2).

Together, these findings demonstrate that *Nf1/Kif1bβ*-deficient tumors reactivate embryonic developmental programs alongside pre-neoplastic and neoplastic cell states, generating spatially organized tumor heterogeneity.

## Discussion

Single-cell and spatial transcriptomic analyses of human neuroblastoma and PPGL reveal shared developmental states of the sympathoadrenal lineage, including early and late embryonic chromaffin-like, connecting progenitor-like (CPC), hybrid, and neuroblast-like identities. These states were spatially organized within tumors, identifying developmental cell-state composition as a source of intratumoral heterogeneity linking these otherwise distinct tumor entities.

Our findings support a developmental model in which neuroblastoma and PPGL arise from a shared sympathoadrenal developmental continuum but diverge through stabilization of different developmental cell states. We propose that these tumors emerge during a developmental window in which immature chromaffin and sympathoblast transcriptional programs remain incompletely fixed and retain the capacity for bidirectional state transitions, such that developmental timing and persistence of plastic intermediate states contribute to tumor phenotype. Developmental single-cell studies established that the fetal sympathoadrenal lineage comprises bridge, connecting progenitor, immature chromaffin, and sympathoblast populations arranged along a common developmental trajectory^7,8^. Subsequent studies have mapped neuroblastoma cells predominantly to fetal sympathoblast states^7,8,10,20^, supporting a developmental origin of neuroblastoma, whereas transcriptional studies of PPGL have largely emphasized chromaffin identity and genetically defined molecular subtypes^13,21^. Whether neuroblastoma and PPGL retain a broader spectrum of sympathoadrenal developmental states beyond their predominant neuroblast- and chromaffin-like identities, and whether such states reflect developmental plasticity, has remained unclear. Here, we show that corresponding chromaffin-, CPC-, hybrid-, and neuroblast-like states occur in both human neuroblastoma and PPGL and are recapitulated during tumorigenesis in genetically engineered mice, providing mechanistic insight into how disrupted sympathoadrenal differentiation alters developmental state transitions.

In wild-type littermates, neuroblast-like states are transient and ultimately resolve toward mature chromaffin identities. In contrast, *Nf1* and *Nf1/Kif1bβ* deficiency biased embryonic differentiation toward neuroblast-like identities and prolonged the persistence of these states before their transition toward a chromaffin fate. Despite apparent restoration of chromaffin identity in the adult adrenal medulla, embryonic developmental programs were reactivated during tumorigenesis, indicating that the capacity to transition between chromaffin and neuroblast states remains latent into adulthood. Rather than enforcing permanent lineage commitment, *Nf1* and *Nf1/Kif1bβ* deficiency appear to preserve the capacity of sympathoadrenal cells to access alternative developmental states. RET provides a potential link between these developmental transitions and neurogenic signaling. *Ret* expression characterized embryonic bridge-neuroblast populations arising from SCP and was high in established tumors, while at the pre-neoplastic stage RET protein was spatially concentrated within cortical breakthrough lesions, where chromaffin-to-neuroblast transitions were observed, but remained low in the surrounding adult adrenal medulla. Given the established neurogenic functions of RET signaling^22,23^, this recurring association with neuroblast-like and transitional states raises the possibility that RET-dependent signaling contributes to the persistence or reacquisition of neurogenic competence. However, whether RET functionally contributes to these transitions, and which extracellular cues and downstream mechanisms instruct neuroblast versus chromaffin fate, remain to be established. The emergence of chromaffin-to-neuroblast transitions within pre-neoplastic lesions prior to overt tumor formation suggests that lineage plasticity is an early event in tumor evolution. These findings extend our previous observation that *Nf1* loss genetically rescues the impaired sympathetic ganglion development caused by *Kif1bβ* deficiency, which disrupts TRKA trafficking and differentiation^17^. The present study links this genetic interaction to developmental plasticity and tumorigenesis and establishes *Kif1bβ* as a tumor suppressor *in vivo*.

These findings provide a potential developmental explanation for the long-recognized occurrence of composite pheochromocytoma–neuroblastic tumors containing chromaffin and neuronal components^24^. Rather than requiring distinct cellular origins, our findings suggest that these components may arise through differential stabilization or reactivation of developmental states within the sympathoadrenal lineage. Rather than representing entirely distinct lineage entities, neuroblastoma and PPGL may therefore be viewed as alternative outcomes of a shared developmental process. Distinct oncogenic drivers likely stabilize different developmental cell states along the sympathoadrenal continuum, thereby shaping tumor phenotype. In this model, neuroblastoma reflects expansion and stabilization of immature neuroblast-like states, whereas PPGL remains enriched for chromaffin-like identities while retaining the capacity to reactivate neurogenic developmental programs. Consistent with this interpretation, a longitudinal MYCN-amplified high-risk neuroblastoma analyzed in this study transitioned from a predominantly neuroblast-like state at initial resection to a chromaffin-dominated state at a second surgery performed six months later. Despite its high-risk molecular features, this patient subsequently became a long-term survivor, raising the possibility that progression toward chromaffin differentiation may be associated with a more favorable clinical course. Although larger cohorts will be required to evaluate this hypothesis, the observation is consistent with a model in which developmental cell state influences tumor behavior beyond conventional diagnostic classification.

Particularly intriguing was the presence of CPC-like, cycling neuroblast, and hybrid developmental states in metastatic PPGLs that were largely absent from non-metastatic tumors. Although late chromaffin-like cells remained the predominant developmental state in both groups, their relative abundance was significantly greater in non-metastatic PPGLs. Together with the chromaffin-to-neuroblast transitions observed in our mouse models, these findings raise the possibility that metastatic progression is associated with a shift away from differentiated chromaffin identity toward more plastic, immature and neuroblast-like developmental states. Although larger cohorts and functional studies will be required to determine whether these states directly contribute to metastatic dissemination, developmental cell-state composition may represent a biologically meaningful dimension of PPGL aggressiveness that complements existing genetic classifications.

Our findings also have implications for tumor classification and therapeutic stratification. Current classification schemes largely separate neuroblastoma and PPGL into distinct diagnostic entities. However, the presence of shared developmental cell states across both tumor types suggests that developmental identity may be as important as histological classification for understanding tumor behavior. This perspective raises the possibility that subsets of PPGL enriched for neuroblast-like programs may share biological vulnerabilities with neuroblastoma, while neuroblastomas containing chromaffin-like populations may retain features of chromaffin differentiation. Although functional studies will be required to evaluate this hypothesis, developmental cell-state profiling may ultimately help identify PPGL patients who could benefit from therapeutic strategies currently used in neuroblastoma.

In summary, our findings demonstrate that differential stabilization and reactivation of developmental programs established during embryonic sympathoadrenal development contribute to heterogeneity within sympathoadrenal tumors. By combining human single-cell and spatial transcriptomics with genetic mouse models, we identify chromaffin-neuroblast plasticity as a developmental mechanism linking neuroblastoma, PPGL and composite tumors, whereby reactivation of embryonic cell states enables tumors to access alternative sympathoadrenal identities.

## Resource availability

### Lead contact

Requests for further information and resources should be directed to and will be fulfilled by the lead contact, Susanne Schlisio.

### Materials availability

This study did not generate new unique reagents. Mouse models used in this study (including Nf1- and Kif1bβ-targeted alleles driven by DBH-Cre) are available from the lead contact upon reasonable request and may require a material transfer agreement (MTA). All other materials generated in this study are available from the lead contact without restriction.

### Data and code availability

Single-cell RNA-seq and spatial transcriptomics data generated in this study have been deposited in and will be publicly available as of the date of publication. Accession numbers will be provided prior to publication. De-identified human tumor data derived from patient sample have been deposited in a controlled-access repository and are available upon request, subject to ethical approval and data access agreements in accordance with Swedish and EU data protection regulations. Summary-level and processed datasets supporting the conclusions of this study will be publicly available at the time of publication. Previously published datasets analyzed in this study are available from the original publications (e.g., Jansky et al., 2021) and relevant public repositories. This paper does not report original code. Any additional information required to reanalyze the data reported in this paper is available from the lead contact upon request.

## Acknowledgments

Funding: W.L. and J.Z were funded by the China Scholarship Council. PB and VPo were funded by Swedish Cancer Fund. M.P. was funded by Swedish Childhood Cancer Foundation. O.C.B.-R. was funded by the Swedish Childhood Cancer Foundation, grants TJ2021-0137 and PR2021-0129. S.S. was funded by the Swedish Research Council, Swedish Childhood Cancer Fund, the Swedish Cancer Society, Knut and Alice Wallenberg Foundation and ParaDifference foundation and supported by ERC Synergy grant (KILL-OR-DIFFERENTIAT). Part of the computations and data handling were enabled by resources provided by the National Academic Infrastructure for Supercomputing in Sweden (NAISS), partially funded by the Swedish Research Council through grant agreement no. 2022-06725. We acknowledge the in situ sequencing facility unit at Science for Life Laboratory for technical assistance with the ISS experiments.

## Author Contributions

Conceptualization: S.S.; Methodology and investigation: W.L., J.Z., V.Pa., P.C., V.Po., M.A., P.B., M.P., J.S., K.S.H., C.L., O.C.B.-R., M.M., I.A., C.C.J., A.T. and S.S.; Computational investigation and analysis: J.Z., M.M., V.Po., M.A., J.S. and O.C.B.-R., Human PPGL sample collection: A.T., C.L. and C.C.J. Human NB sample collection: P.K. and J.S. Mouse adrenal tumor diagnosis: C.C.J. and A.T. Writing, S.S., A.T. and J.Z. Funding Acquisition, Resources & Supervision, S.S., Competing interests: none.

## Declaration of generative AI and AI-assisted technologies

Portions of the manuscript text were edited with the assistance of a large language model (ChatGPT, OpenAI) to improve the readability and language of the manuscript. All scientific content, interpretations, and conclusions were generated and verified by the authors, who reviewed and edited the final manuscript and take full responsibility for its content. After using this tool, the authors reviewed and edited the content as needed and take full responsibility for the content of the publication.

## Supplemental information titles and legends

Supplementary Table 1-2 and Supplementary Data 1-12

## METHODS

### Human single-nucleus RNA sequencing

Human single-nucleus RNA sequencing experiments were performed following the protocols described in our previous studies^12,15^, including sample preparation, library construction, sequencing, quality control, and downstream analyses.

### Reference-based label transfer and hybrid-state annotation

Single-nucleus RNA sequencing tumor datasets were reprocessed from raw count matrices for label-transfer analysis. Raw counts were normalized using the SCTransform method implemented in Seurat (reference to Seurat, version will be after server is back online) and the 2,000 most highly variable features were selected for downstream analyses. Principal component analysis was computed on the normalized expression matrix.

Label transfer was then performed using the FindTransferAnchors and TransferData functions in Seurat (reference to Seurat, versions will be written when server is on again). A healthy adrenal gland reference atlas from Jansky et al. ^25^ served as the reference dataset, and healthy cell state annotations were assigned to tumor cells based on shared transcriptional profiles identified through anchor mapping. Because cancer cells are not expected to correspond one-to-one with healthy cell identities, transferred labels were interpreted as measures of transcriptional similarity rather than definitive cell-type assignments.

For each tumor cell, label transfer scores were extracted for all reference cell-type populations. Cells with a maximum label-transfer score below 0.25 were classified as low-confidence and excluded from subsequent analyses. These cells were considered insufficiently similar to any healthy reference population and may represent tumor-specific, transitional, out-of-distribution, or otherwise poorly classified transcriptional states. For the remaining cells, competing reference identities were evaluated relative to the highest-scoring label for that cell. Any reference label with prediction scores equal to or greater than 80% of the highest-scoring label for that cell was retained. Cells retaining a single label were classified as clear assignments and correspondingly got labeled with that reference cell type. Cells retaining multiple labels were classified as hybrid-like states, indicating substantial transcriptional similarity to multiple healthy reference populations. Hybrid labels were generated by concatenating all retained reference identities into a composite label. Importantly, these hybrid-like annotations were not interpreted as evidence for the existence of true biological hybrid cell types. Rather, they provide a practical framework for identifying tumor cells whose transcriptional profiles resemble multiple healthy cellular states while distinguishing these from cells that show limited resemblance to any reference population.

### Bulk RNA-seq data deconvolution

Bulk RNA-seq data and matched clinical metadata were obtained from a previously published human PPGL cohort (hereafter referred to as the CNIO cohort) ^16^. Cell-state composition of the CNIO cohort was estimated using the Bayesian deconvolution framework BayesPrism^26^. A single-cell reference dataset was constructed from the PPGL single-nucleus RNA-seq data generated in this study, including both tumor and stromal cell populations. Tumor cells were annotated using the reference-based label-transfer approach described above, and the resulting annotations were used to define tumor cell states. Healthy adrenal chromaffin cells were included as an additional reference cell population.

Prior to deconvolution, genes shared between the single-cell reference and bulk RNA-seq datasets were retained, and raw count matrices from both datasets were used as input. Cell states represented by fewer than 20 reference cells were excluded from the reference dataset. BayesPrism was run using the default workflow and parameters to estimate the contribution of each reference cell state to the bulk transcriptomic profiles. Deconvolution results were reported as BayesPrism θ estimates, representing the proportion of total RNA contributed by each reference cell state within each bulk RNA-seq sample.

### Visium HD Spatial Transcriptomics Library Construction, Sequencing and Preprocessing

cDNA library of the human pheochromocytoma Formalin-Fixed Paraffin-Embedded (FFPE) tissue was generated using the Visium HD Spatial Gene Expression Reagent Kits with Human Transcriptome Probe Kit v2 (10x Genomics), following the manufacturer’s instructions. The resulting library was then sequenced on NovaSeq X Plus 25B-300 flow cell (NGI, Stockholm, Sweden). Raw sequencing data were demultiplexed and subjected to quality control by the sequencing facility and delivered via the NGI Data Delivery System (DDS). Spatial gene expression matrices were generated using Space Ranger v4 (10x Genomics). Raw sequencing data were mapped to the human GRCh38 reference genome and the corresponding Human Transcriptome Probe Kit v2 probe set sequences. Cell segmentation was performed using the built-in nucleus-based segmentation module by enabling the --nucleus-segmentation option (--nucleus-segmentation=true), with a maximum nucleus diameter of 50 pixels (--max-nucleus-diameter-px=50) and a nucleus expansion distance of 4 μm (--nucleus-expansion-distance-micron=4), resulting in cell-level expression profiles.

### Visium HD Spatial Transcriptomics Data Analysis

Cell-gene count matrix was analyzed by using Seurat ^27^. Cells with number of detected genes lower than 20 or percentage of mitochondrial gene higher than 20 were excluded. Then, the gene count matrix was normalized and variance-stabilized using SCTransform, with a subsampling of 10,000 cells. PCA was then performed on the normalized data. The top 20 principal components were used to construct a shared nearest neighbor (SNN) graph and for Uniform Manifold Approximation and Projection (UMAP) low-dimensional visualization. Cell clustering was performed using the Louvain algorithm at a resolution of 0.6. Differentially expressed gene analysis was conducted by using the FindAllMarkers function with the Wilcox test. Cell clusters were annotated based on the expression level of canonical cell type marker genes. The neuroendocrine cell populations were then projected onto cell states defined in an embryonic adrenal medulla single-cell dataset from Jansky et al. ^25^ using the Seurat-based label-transfer procedure described above. Final cell-state annotations were assigned based on prediction scores, using an absolute margin of 0.2 from the highest prediction score to identify competing reference cell-state assignments. Cells with a maximum prediction score below 0.4 were classified as low-confidence. Cells with multiple competing assignments were annotated as hybrid-like states, whereas cells with a single assignment were labeled according to the highest-scoring reference cell state.

### Cell-type and cell-state kernel density estimation

Spatial enrichment of selected cell populations was visualized using two-dimensional kernel density estimation based on the spatial coordinates of individual cells. Kernel density was estimated using the kde2d function implemented in the MASS R package and the estimated density values were mapped back to the corresponding cell coordinates for visualization

### Mouse lines

Mice experiments were performed in accordance with the ethical permit issued by the Ethical Committee on Animal Experiments and in accordance with the Swedish Animal Agency’s Provisions and Guidelines for Animal Experimentation recommendations. R26RTomato, R26RYFP, and *Sox10*CreERT2 mice were received from the Adameyko lab (Karolinska Institutet). *KIF1B*βfl/fl; *Dbh*Cre mice are described in Li et al. (2016) and *NF1*fl/fl mice were described recently in Zhu et al. ^19^ and were obtained from the Jackson laboratory [mouse strain B6.129(Cg)-*Nf1*tm1par/J].

Timed mattings were started in the afternoon. The day of plug detection was defined as E0.5 and the day of birth as P0. For the analysis of embryonic stages, oral gavage of the plug-positive female mouse was performed with 2 mg of tamoxifen (Sigma, T5648, dissolved in corn oil) at the specified embryonic stage (E8.5 or E11.5). For postnatal induction of Cre expression, oral gavage of the nursing mouse was performed daily with 2 mg tamoxifen for 5 days (P0-4).

### Sample collection and single cell sequencing

Forty mouse adrenal anlagen/gland samples with different genotypes and developmental stages (detailed in Supplementary Data 1) were collected via surgical resection after perfusion. For embryonic tissue collection, the pregnant mouse was euthanized by carbon dioxide inhalation on an embryonic day 17 (E17.5). The embryos were collected, decapitated and the body was transferred into ice-cold 1x PBS (for immunohistochemistry and RNAscope analysis or LBSS (Locke’s balanced salt solution, for subsequent single-cell RNA-seq analysis). Embryonic adrenal glands were isolated under the dissection microscope. For postnatal tissue isolation, mice older than 13 days were euthanized by carbon dioxide inhalation. After euthanasia, adrenal glands were immediately dissected.

The tissue from each sample was dissociated and sorted in 384-well plates. Sorted cells were lysed and their RNA was obtained using the Smart-seq2 protocol. Libraries were prepared using the Tn5 transposase tagmentation (Nextera XT), and their quality was assessed with fragment analyzer. High-quality libraries were sequenced using Illumina HiSeq 2500, and further de-multiplexed with deindexer (https://github.com/ws6/deindexer) using the Nextera index adapters and the 384-well layout. Forty-two libraries for the forty adrenal samples adding up 16,128 cells were sequenced, producing in total 9.91x10^9^ single-end reads.

### Sample preparation for single-cell sequencing

For single-cell suspension preparation for single-cell RNA-seq, multiple adrenal glands from pups within one litter were pooled in 1 ml LBSS (154 mM NaCl, 5 mM KCl, 3.6 mM NaHCO3, 5 mM HEPES, 11 mM glucose) on ice after cutting them open to expose the medulla. The cortex from adrenal glands of mice older than 13 days was partially peeled off to enrich for medullary cells. For mice older than 3 months, adrenal glands/tumors were collected after avertin anesthesia and 30ml LBSS perfusion.

After LBSS was exchanged 3 times, the tissue was incubated under occasional agitation in 500 µl pre-warmed papain solution (25 U/ml) in dissociation medium (DMEM/F12 medium without dye with HEPES buffer supplemented with 1mM L-Cysteine, 1mM CaCl2, 1mM EDTA) for 30 min at 37°C. Afterward, the medium was replaced 3x by 1 ml trituration buffer (DMEM/F12 medium without dye with HEPES buffer supplemented with 2% Horse Serum, 1mM CaCl2, 1mM EDTA) and eventually by 500 µl trituration medium and 12.5 µl RNAsin and triturated gently 20x with 3 fire-polished Pasteur pipettes of decreasing diameter.

Subsequently, the cell suspension was filtered through a 40 µm strainer and single cells were sorted into 384 well plates prefilled with Smart-seq2 lysis buffer. Plates were immediately sealed and transferred to dry ice to the Eukaryotic Single Cell Genomics (ESCG) facility at SciLifeLab for further processing.

### Read mapping, quality control, and gene expression calculation

To select high quality reads we allowed for a hard-clipping of adaptors (using CutAdapt 1.13), allowing for a maximum error rate of 0.1 and excluding reads with less than 20 bases. Additional diagnostics on the reads quality were conducted with FASTAQC (https://www.bioinformatics.babraham.ac.uk/projects/fastqc), cells with reads failing at three or more quality control (QC) tests (among eleven in total including per base- and sequence quality scores, frequency of each nucleotide, GC content, over-represented motifs, and number of duplicated reads) were excluded for further analysis.

High quality reads were mapped with STAR^28^ using 2-pass alignment to have improved performance of *de novo* splice junction reads, to the mm10 mouse genome version mm10/GRCm38.p6 (released 2019), annotated by the comprehensive annotation of GENCODE 18 ^29^ obtained from the UCSC browser ^30^ on 14.01.2019. Alternative chromosomes were excluded from the annotations. Gene expression was calculated using HTSeq ^31^.

To conduct a QC of the cells, different technical (i.e. proportion of mapped reads, multi-mapped reads proportion, not aligned reads proportion, number of detected genes, transcriptome variance, # very highly expresed genes, # highly expressed genes, # moderatly expressed, # lowly expresed, # very lowly expressed, Cell-to-Mean-Correlation, and Number of highly expressed and highly variable genes) and biological features (i.e. aptopotic proportion, mtDNA proportion, metabolic proportion, ribosome proportion, Actb, Gadph, membrane proportion, cytoplasm proportion, extracellular region proportion, mitochondria upregulated in broken cells, mitochondria downregulated in broken cells) were calculated with QoRT ^32^ and Celloline ^33^. Using these features, cells with low quality were identified using Cellity ^33^. Briefly, Cellity runs an unsupervised classification of cells based on the most relevant features over a PCA to differentiate cells of good- from those of poor-quality. Cells with good quality from the analysis of merged cells from all samples (i.e. E17 and later stages) were selected for further analysis.

### Cell clustering and expression analysis

For cell clustering analysis, samples were processed in age batches (i.e. E17 and later groups independently). Gene expression of cells/nuclei was filtered, transformed, scaled and standardized accounting for sequence depth, mitochondrial content, and cell-cycle stage, using Scanpy ^34^. After manual inspection, cells with at most 10% of mitochondrial genes, and a minimum number of 200 genes and at most 10,000 were analyzed. Genes expressed in less than 3 cells were further excluded from the analysis. 6,274 cells for 29 libraries and 27 samples adding up 2.52x10^9^ single-end reads were included in the post-natal adrenal analysis; and 4,166 cells for 13 samples/libraries adding up 1.50x10^9^ single-end reads in the E17 adrenal analysis. Counts for the selected genes and cells were normalized as the log(reads per 10,000 + 1). To account for depth and mitochondrial content, both the total read counts and percentage of mitochondrial genes per cell were regressed.

The processed counts were scaled to unit variance and zero mean, filtering out genes exceeding a standard deviation of 10. Cell-cycle effect was corrected by regressing the mouse orthologue cell-cycle genes to the human set reported by Tirosh et al. ^35^. With the resulting expression values, a graph of nearest neighbor cells was computed using the first 40 component from a PCA, and 10 neighbor nodes, and clustered with the Louvain algorithm^36^.

The normalized gene expression (i.e. log[reads per 10,000 + 1]) was used to determine the genes significantly up-regulated in each cluster (i.e. genes whose average expression is significantly higher than the average expression of cells in all the other clusters). This significance was calculated using Benjamini-Hochberg multiple test correction on Welch’s *t*-test generated *p*-values, with a FDR threshold of 0.01.

### Cell velocities and differentiation potential analysis

To calculate the differentiation trajectory of cells, the percentage of splicing and unspliced genes per cell was estimated with velocyto [10]. Further, the cell velocities were calculated using scVelo ^37,38^. To obtain high confidence velocities (as confirmed by visual inspection of confidence scores) we selected genes with a minimum number of shared read counts of 2,500, and built the velocities using the top 2,000 genes. First-and second-order moments for velocities estimation were computed for each cell across its nearest 10 neighbors from a neighbor graph obtained from euclidean distances in the top 50 components from a PCA.

Further, the gene transition over the velocity pseudotime (representing random-walk based distance measures on the velocity graph, and the approximate time of cell differentiation) was determined using a stochastic model, and illustrated for the selected populations of interest. The differentiation potential of cells was determined with Palantir ^39^ using the default parameters, and as starting cell the one with the highest expression of the precursor marker *Erbb3*. This approach uses entropy to estimate cell plasticity in modeled trajectories of cells fates, so that cell plasticity increases with entropy.

### Gene regulatory network reconstruction

Gene regulatory networks were inferred using the pySCENIC (v0.12.1) workflow. Co-expression modules were first identified using gradient boosting–based regression (GRNBoost) with a curated list of mouse transcription factors. This step resulted in an adjacency matrix representing putative transcription factor–target gene relationships.

To refine co-expression modules into high-confidence regulons, cis-regulatory motif enrichment analysis was performed using mouse cisTarget reference databases. Motif enrichment was assessed in both promoter-proximal and distal regulatory regions using the following databases: RefSeq-based rankings covering ±10 kb around transcription starts sites and −500 bp upstream to +100 bp downstream of transcription start sites. Motif annotations were obtained from the non-redundant mouse motif collection (motifs-v9-nr, MGI).

Refined regulons were subsequently used to compute per-cell regulon activity scores using AUCell, yielding a regulon activity matrix that captures the activation state of transcriptional regulatory programs across individual cells.

### Identification of overdispersed genes and low-dimensional embedding

Highly overdispersed genes were identified using scFates variance modeling. Gene expression values were scaled with clipping at a maximum value of 10 to reduce the influence of extreme outliers. Principal component analysis (PCA) was performed on the scaled expression matrix restricted to overdispersed genes.

A k-nearest neighbor graph was constructed in PCA space using cosine distance across the first 30 principal components. Low-dimensional visualization was generated using UMAP, and unsupervised cell clustering was performed using the Leiden community detection algorithm.

### Diffusion map construction and manifold refinement

Diffusion maps were computed using Palantir on the first 50 principal components with a neighborhood size of 80 cells to capture continuous cellular state transitions. The leading diffusion components (DC1–DC8) were retained as a reduced representation of the developmental manifold.

A k-nearest neighbor graph was reconstructed in diffusion space using 40 neighbors and used to infer global connectivity between clusters via Partition-based Graph Abstraction (PAGA). To preserve large-scale lineage topology in low-dimensional visualization, UMAP embeddings were initialized using the PAGA graph structure.

For final embedding refinement, a nearest neighbor graph was recomputed in diffusion space using 80 neighbors, followed by UMAP projection with a spread parameter of 0.6. Regulon activity scores were integrated into an AnnData object while preserving diffusion embeddings, UMAP coordinates, and cell-level metadata. This enabled trajectory inference to be performed directly on transcriptional regulatory program dynamics rather than raw gene expression.

### Principal graph–based trajectory inference and dynamic regulon analysis

Trajectory inference was performed using scFates (v1.0.9) employing the SimplePPT approach on the diffusion embedding. For each dataset, a principal graph consisting of 2000, 1200, and 900 nodes was fitted using dataset-specific hyperparameters (scFates tl.tree, method = “ppt”, sigma = 002/002/008, lambda = 1100/1100/1600, metric = “euclidean”). To improve biological interpretability, short spurious branches were pruned from the inferred trees. Root nodes were selected based on early developmental positioning within the diffusion manifold. Pseudotime values were computed as geodesic distances along the principal graph and projected onto individual cells using soft assignment weights. Dynamic changes in transcriptional regulatory programs along developmental trajectories were quantified using generalized additive models implemented in scFates. Spline-based regression was applied to identify regulons exhibiting significant associations with pseudotime. Fitted smooth curves were subsequently used to model continuous activation and repression patterns of regulatory programs across lineage progressions.

### In-Situ Sequencing

The mouse tissue sections were processed by *in situ* sequencing following the In-Situ Sequencing (ISS) protocol ^40^. We selected 100 gene-of-interest for the purpose of ISS. The full list of gene-of-interest can be found in Supplementary Data 9.

### In-Situ Sequencing Data Preprocess

The ISS data from all mouse tissue sections were first processed manually. Briefly, RNA signals were projected onto the corresponding DAPI staining images using TissUUmaps ^41^. Tissue regions were manually delineated based on the nuclear staining, and signals located outside the tissue boundaries were excluded. Tissue sections were hexagonally binned with a radius of 250 pixels (approximately 40 μm) using the sf R package ^42,43^. The bin centers were used as spatial coordinates, and RNA signals falling within the same bin were aggregated. This resulted in a pseudo-spot spatial gene expression matrix for each tissue section.

### Pseudo-spot Spatial Transcriptome Analysis

The pseudo-spot spatial transcriptomic data were further analyzed using Giotto ^44^. Briefly, the pseudo-spot spatial gene expression matrices were used as input. A gene was considered expressed if its count was greater than or equal to 1, and genes detected in fewer than 5 pseudo-spots were removed. Pseudo-spots expressing fewer than 5 genes were regarded as low quality and excluded from downstream analyses. The raw gene count matrices were normalized using the normalizeGiotto function with default parameters. Principal component analysis (PCA) was then performed using runPCA. A nearest-neighbor (NN) network was constructed using createNearestNetwork based on the top 30 PCs, with 20 nearest neighbors identified for each hexagonal bin. Clustering was performed using the doLeidenCluster function, which uses the Leiden community detection algorithm with a resolution parameter of 0.5.

### Deconvolution of Pseudo-spot Spatial Transcriptome

Mouse single-cell RNA-Seq data in this study were used as reference, and Robust Cell Type Decomposition (RCTD) ^45^ was applied to infer the cellular composition of the hexagonally binned spatial transcriptomic data. The reference object was constructed using the Reference function from the spacexr R package ^45^, with the raw count matrix of non-immune cells as input. A SpatialRNA object was generated using the spatial coordinates and gene expression profiles of the bins. The RCTD object was created using create.RCTD, with both UMI_min and counts_MIN set to 5. Deconvolution was performed using run.RCTD. Predicted normalized cell type weights were extracted as the final deconvolution results.

### Immunohistochemistry

For OCT-embedded tissue, dissected tissue was transferred to 4% paraformaldehyde solution for 24h at 4 degrees. After washing with PBS, they were incubated in 10% sucrose and overnight in 30% sucrose. Afterward, the tissue was transferred into OCT and stored at −80 °C. 10 µm sections were prepared and collected on adhesives slides. After drying for 15 min, the slides were washed 3x 5 min in PBS. Afterward, the tissue was incubated in a blocking solution (1x PBS with 0.1% triton, 5% horse serum) for 60 minutes. Afterward, sections were incubated with the respective antibodies and concentrations in blocking solution overnight at 4°C. After 3x 5 min PBS washing steps, the tissue was incubated with Alexa647/555/488 antibodies (1:1000 in blocking solution) for 2h at RT protected from light. After 3x 5 min PBS washing steps, sections were incubated for 30 sec with the autofluorescence quencher true black (1:20 in 70% ethanol). Subsequently, the sections were washed 5x 5 min with PBS, incubated with 1 µg/ml DAPI for 5 min, and mounted with ProLong™ Gold Antifade Mountant with DAPI.

For 4% paraformaldehyde-fixed paraffin-embedded (FFPE) tissue, tissue was fixed in 4% paraformaldehyde solution at 4 °C for 24h. After washing with PBS, tissue was transferred to an automated tissue processor and embedded with paraffin. 4um sections were collected and stored at 4 °C. Before staining, paraffin sections were dried at 60 °C for 1h, followed by deparaffination at xylene and rehydration in different concentrations of ethanol, and finally immersed in water. Samples were subjected to antigen retrieval with 1× Dako Target Retrieval solution (Dako, 1699). The staining protocol is the same as OCT embedded tissue.

### Hematoxylin and eosin H&E staining

4um FFPE sections were stained using the Hematoxylin and Eosin Stain Kit (Vector Laboratories, H-3502), 1min of hematoxylin, and 3 min of eosin were used according to the manufacturer’s instructions. Slides were mounted with Mounting agent Pertex (Histolab, 00840-05) and dried in a fume hood until imaging.

### Primary antibodies

Goat anti-PHOX2B (1:40, R&D Systems, AF4940), chicken anti-NEFH (1:2000, Abcam, ab4680), rabbit anti-CHGB (1:500, Synaptic Systems, 259 103), sheep anti-TH (1:1000, Novus Biologicals, NB300-110), were used and diluted in blocking solution and applied on sections overnight at 4°C.

### Fluorescent RNA *in situ* hybridization (RNAscope)

After dissection, left adrenal glands were snap-frozen on dry ice and stored at −80 °C. The native tissue was mounted on a drop of OCT in the cryotome before sectioning. 10 µm sections were collected on adhesive slides and after 1 h incubation at −20 °C were stored at −80 °C.

RNA *in situ* hybridization was performed using the RNAscope® Multiplex Fluorescent Detection Kit v2 on native sections after 1 h post-fixation in 4% PFA at RT, ethanol dehydration steps, and 10 min protease IV treatment at RT according to the manufacturer’s instructions for native/FFPE tissue.

The human-specific probes used are *Nefm* (575251), *Chgb* (567681-C2), *Mki67* (591771), and *Stmn4* (850151). The mouse-specific probes used are *Nefm* (315611), *Chgb* (1057371-C2), *Th* (317621-C3), *Mki67* (416771), *Prph* (400361-C3), *Isl1* (451931), *Hand1* (429651), *Pnmt* (426421-C3), *Stmn4* (537541), *Gap43* (318621-C3), *Nefl* (493671), *Ret* (431798-C2), *Alk* (501131), *Mycn* (477151-C3).

Positive cells in medullary area were counted by using Cellpose-SAM and ImageJ (Pachitariu, M., Rariden, M., & Stringer, C. (2025). Cellpose-SAM: superhuman generalization for cellular segmentation. bioRxiv.). For the averaged percentages of medullary areas (TH^+^) over total adrenals (DAPI^+^) and *Nefm*^+^ cells over total medullary (DAPI^+^) cell number, the Shapiro–Wilk test and Brown–Forsythe test were first used to assess normality and homogeneity of variances. Since the data satisfied the assumptions of normality and homogeneity of variances, one-way ANOVA followed by Tukey’s post hoc test was used. Otherwise, the Kruskal–Wallis test followed by Dunn’s post hoc test was applied, and the corresponding *p*-values are reported. *P*-values < 0.05 were considered statistically significant. Data are presented as mean ± s.e.m. Note that *n* in all experiments represents the number of biologically independent samples (individual mouse adrenal).

### Microscopy

Images acquisition was performed with LSM800 Zeiss confocal microscope and Zeiss AxioScan.Z1 Slide Scanner using a 20x objective. They were acquired as .czi files and processed with ZEN 3.0(blue edition).

### Ethical Considerations

Human pheochromocytoma tumor#1 was obtained and studied with patients’ informed consent and approval by the Tufts Medical Center Institutional Review Board.

Collection and analyses of all other human PPGL samples were obtained with patients’ informed consent and approval covered by the ethical approval numbers 01-136 local ethical committee KI forskningsetikkommitté Nord) and 2020-04226 (Swedish Ethical

Review Authority). Post-mortem human adrenal glands for staining’s were obtained from the NIH Neurobiobank (University of Maryland, MD) under the same ethical permit from Stockholm Regional Ethical Review Board and the Karolinska University Hospital Research Ethics Committee (KI 2007/069 and KI 2001/136). Ethical permits for neuroblastoma samples (009/1369-31/1 and 03-736) were issued to Professor P. Kogner. Neuroblastoma tumor samples were collected after obtaining verbal or written consent from parents or guardians depending on IRB requirements at the time of sampling. All samples were collected and analyzed according to permits approved by the Karolinska Institutet and the Karolinska University Hospital ethics committees (reference numbers 03-736 and 2009/1369-31/1) in agreement with the Declaration of Helsinki. All tumors sampling was performed during clinical routine procedures, treatment and management according to established national and international protocols, and hospital records were used to retrieve clinical data.

Ethical permits for animal studies were approved by the appropriate local and national authorities Jordbruksverket, Sweden.

## CONTACT FOR REAGENT AND RESOURCE SHARING

Further information and requests for resources and reagents should be directed to and will be fulfilled by the Lead Contact, Susanne Schlisio

## Figure legends

**Extended Data 1.**
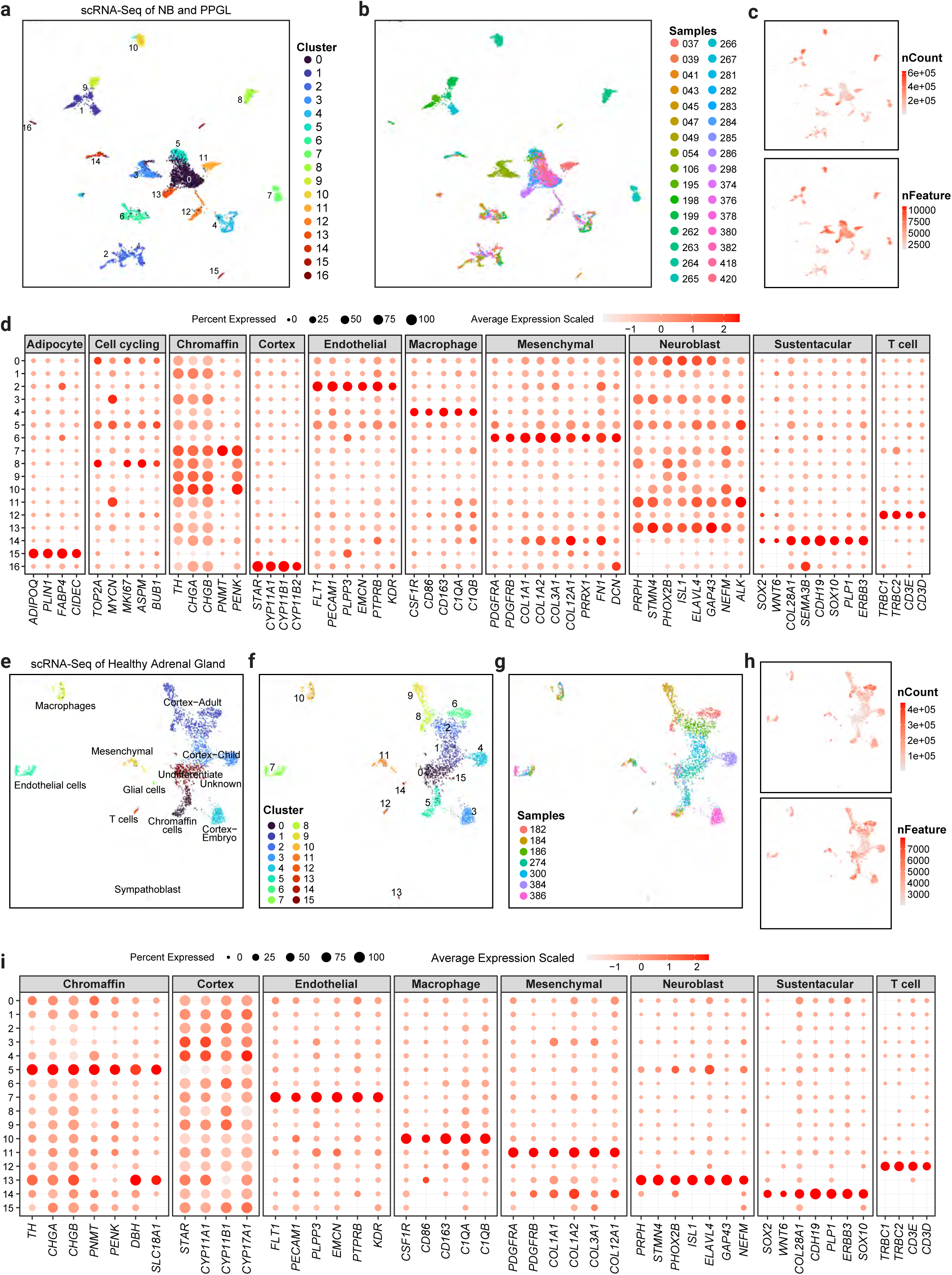
Quality control and annotation of PPGL, NB and healthy adrenal gland single-nucleus datasets. **a-d** Quality control and annotation of the merged PPGL–NB single-nucleus RNA-sequencing dataset. (a) UMAP of the merged dataset colored by unsupervised clusters. **b** UMAP colored by sample identity. **c** UMAPs showing total UMI counts (*nCount*, upper panel) and the number of detected genes (*nFeature*, lower panel) for each cell. **d** Dot plot showing canonical marker genes used to annotate tumor and non-tumor cell populations. Dot color indicates scaled average expression and dot size indicates the percentage of cells expressing each gene. **e-i** Quality control and annotation of the healthy human adrenal gland Smart-seq2 reference dataset. **e** UMAP colored by annotated cell type. **f** UMAP colored by unsupervised clusters. **g** UMAP colored by sample identity. **h** UMAPs showing total UMI counts (*nCount*, upper panel) and the number of detected genes (*nFeature*, lower panel). **i** Dot plot showing canonical marker genes used to annotate the healthy postnatal adrenal cell populations.

**Extended Data 2.**
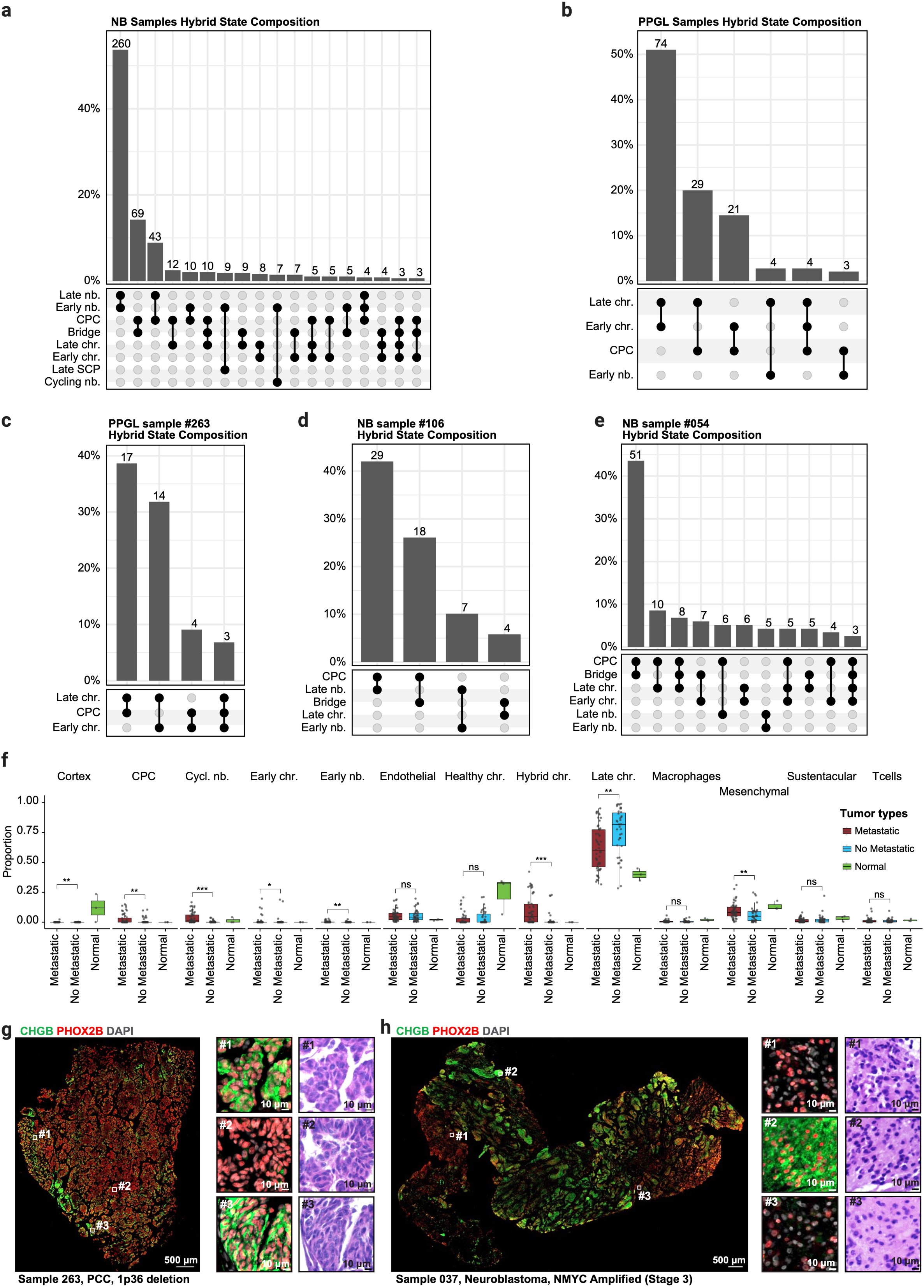
Hybrid developmental labels and extended deconvolution analyses. **a-e** UpSet plots summarizing hybrid developmental-state assignments following label transfer from the human fetal adrenal medullary reference atlas. Bar heights indicate the number of tumor cells assigned to each hybrid label combination, and the connected dots indicate the contributing developmental states. **a** Hybrid developmental-label combinations identified across all NB tumor cells. **b** Hybrid developmental-label combinations identified across all PPGL tumor cells. **c** Hybrid developmental-label combinations in PPGL sample #263. **d** Hybrid developmental-label combinations in NB sample #106. **e** Hybrid developmental-label combinations in NB sample #054. **f** Comparison of inferred cell-state proportions across metastatic PPGL, non-metastatic PPGL and normal adrenal medulla for all deconvoluted cell states. Box plots show the median and interquartile range, with points representing individual samples. Statistical significance was assessed using two-sided Wilcoxon rank-sum tests comparing metastatic and non-metastatic PPGLs, followed by Benjamini–Hochberg false-discovery-rate (FDR) correction. ns, FDR ≥ 0.05; *FDR < 0.05; **FDR < 0.01; ***FDR < 0.001. **g,h** Representative multiplex immunofluorescence staining of CHGB (green), PHOX2B (red) and DAPI (gray) in PPGL sample #263 (PCC with chromosome 1p36 deletion) (g) and MYCN-amplified stage 3 neuroblastoma sample #037 (h). White boxes indicate regions shown at higher magnification, alongside the corresponding H&E-stained sections. Scale bars, 500 μm (whole-section images) and 10 μm (higher-magnification images).

**Extended Data 3.**
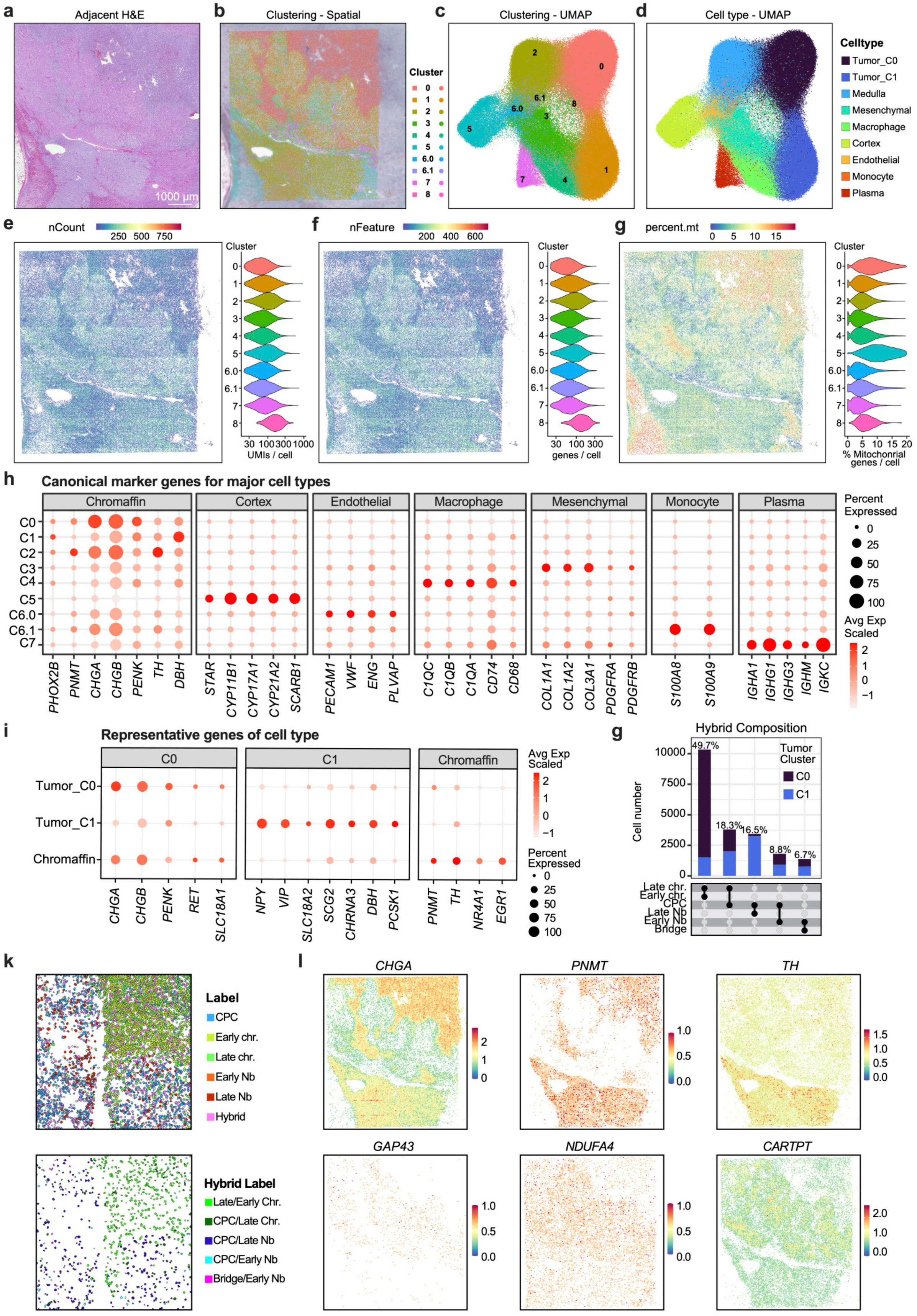
Quality control, clustering and cell-type annotation of PPGL #1 Visium HD spatial transcriptomic data. **a** H&E-stained adjacent tissue section corresponding to the region analyzed by Visium HD spatial transcriptomics. Scale bar, 1,000 μm. **b** Spatial distribution of unsupervised transcriptional clusters across the tissue section. **c** UMAP of unsupervised transcriptional clusters. **d** UMAP showing annotated tumor and non-tumor cell populations. **e-g** Spatial maps and corresponding violin plots showing post-filtering quality-control metrics, including total UMI counts per cell (**e**), number of detected genes per cell (**f**), and percentage of mitochondrial transcripts (**g**). **h** Dot plot showing canonical marker genes used for annotation of major cell types. Dot color indicates scaled average expression and dot size indicates the percentage of cells expressing each gene. **i** Dot plot showing representative marker genes distinguishing the two tumor clusters (Tumor_C0 and Tumor_C1) and normal chromaffin cells. Dot color indicates scaled average expression and dot size indicates the percentage of cells expressing each gene. **j** UpSet plot summarizing hybrid developmental-state assignments following label transfer from the human fetal adrenal medullary reference atlas. Bar heights indicate the number of tumor cells assigned to each hybrid-state combination, and connected dots indicate the contributing developmental states. **k** Spatial distribution of transferred developmental cell-state labels (upper) and hybrid developmental-state labels (lower) within the ROI. **l** Spatial expression of representative marker genes, including *CHGA*, *PNMT*, *TH*, *GAP43*, *NDUFA4*, and *CARTPT*.

**Extended Data 4.**
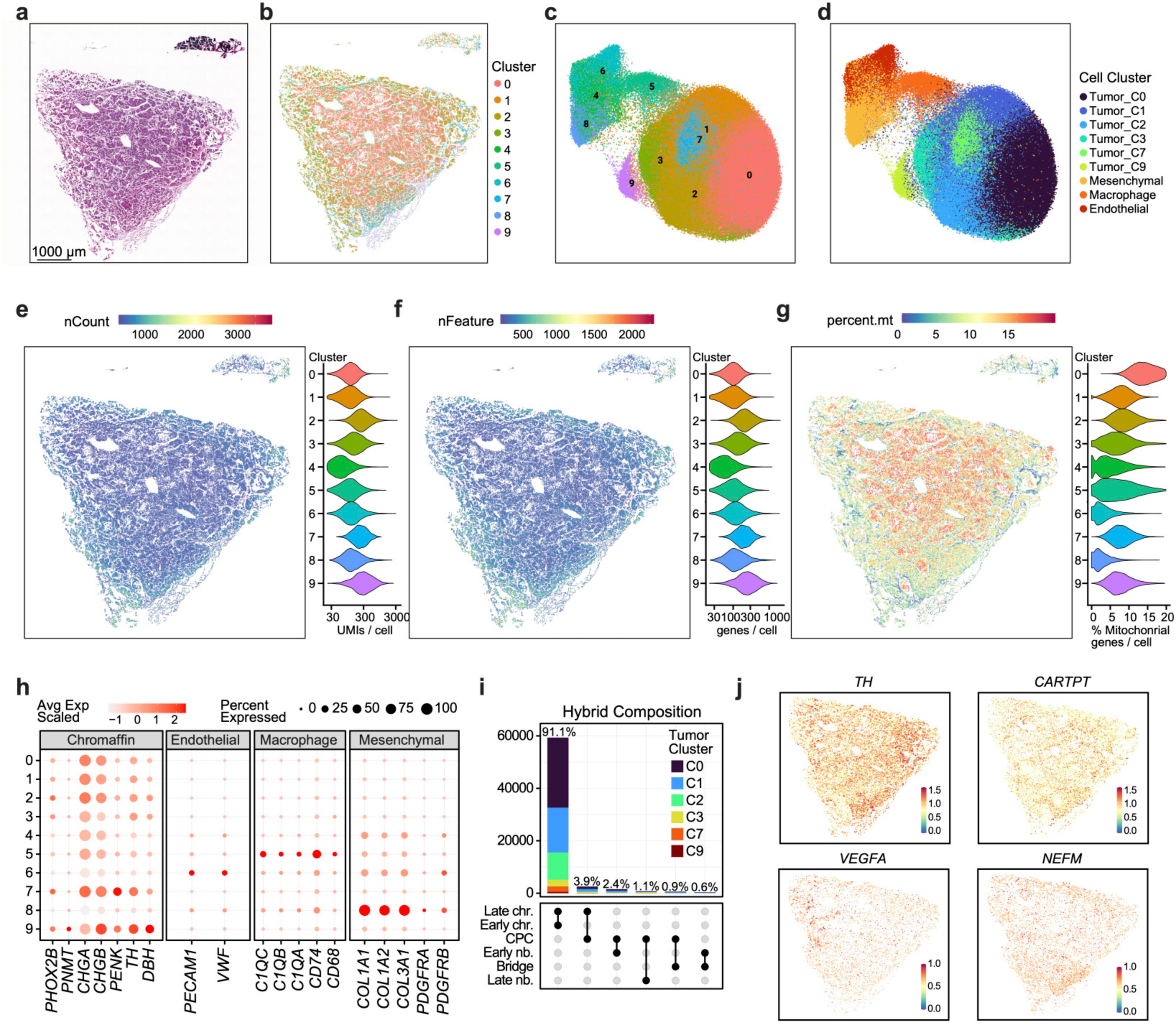
Quality control, clustering, developmental cell-state mapping, and regional heterogeneity in malignant PPGL #263 analyzed by Visium HD. **a** Hematoxylin and eosin (H&E)-stained tissue section corresponding to the region analyzed by Visium HD spatial transcriptomics. Scale bar, 1,000 μm. **b** Spatial distribution of unsupervised transcriptional clusters across the tissue section. **c** UMAP of unsupervised transcriptional clusters. **d** UMAP showing annotated tumor and non-tumor cell populations. **e-g** Spatial maps and corresponding violin plots showing post-filtering quality-control metrics, including total UMI counts per cell (**e**), number of detected genes per cell (**f**), and percentage of mitochondrial transcripts (**g**). **h** Dot plot showing canonical marker genes used for annotation of major cell types. Dot color indicates scaled average expression and dot size indicates the percentage of cells expressing each gene. **i** UpSet plot summarizing hybrid developmental-state assignments following label transfer from the human fetal adrenal medullary reference atlas. Bar heights indicate the number of tumor cells assigned to each hybrid-state combination, and connected dots indicate the contributing developmental states. **j** Spatial expression of representative marker genes, including *TH*, *CARTPT*, *VEGFA*, and *NEFM*.

**Extended Data 5.**
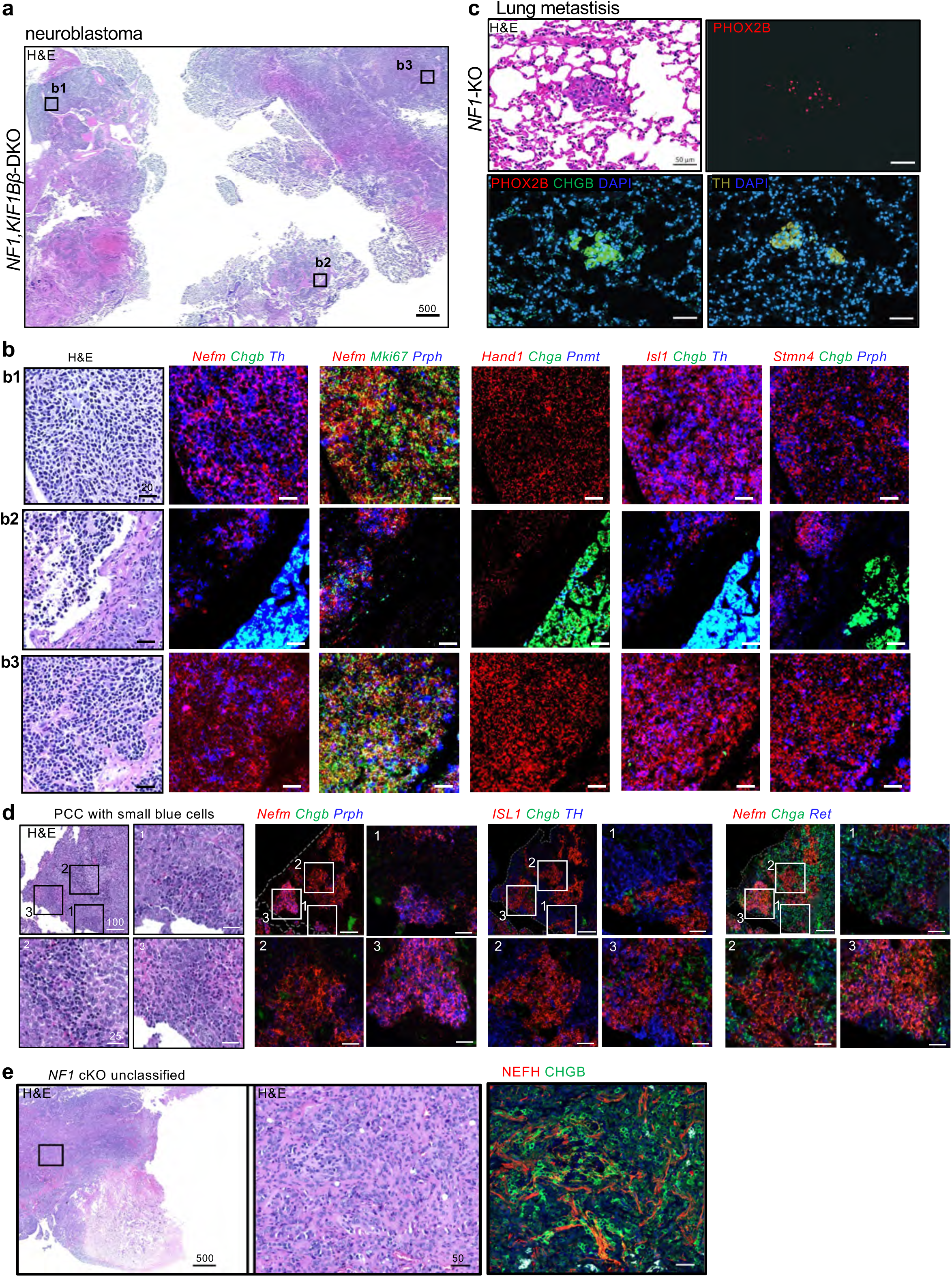
Extended histopathological and molecular characterization of representative tumors. **a-b** Extended RNAscope characterization of the *NF1/KIF1Bβ*-cKO neuroblastoma shown in Fig. 1D. Multiple sections stained for *NEFM, CHGB, PRPH, ISL1, TH*, and additional markers demonstrate predominant expression of neuroblast-associated markers (*NEFM, ISL1, PRPH*), consistent with neuroblast identity. *TH* expression is observed across tumor regions, reflecting catecholaminergic lineage identity. While most tumor areas lack chromaffin markers (*CHGB*), focal regions (e.g., b2) display *CHGB*⁺ cells, indicative of localized chromaffin-like differentiation. Scale bars: overview, 500 μm; inset, 30 μm. **c** Additional lung metastasis from an *NF1*-cKO mouse. H&E shows tumor cell clusters within lung parenchyma. Immunostaining for PHOX2B, CHGB, and TH confirms chromaffin lineage identity of metastatic lesions. Scale bars as indicated 50 μm. **d** Pheochromocytoma with small-cell foci from an *NF1*-cKO mouse (same case as Fig. 1g). H&E shows classic PCC architecture with focal aggregates of small, densely nucleated cells. RNAscope for *Nefm, Chga/Chgb, Ret, Mki67,* and *Prph* demonstrate that the small-cell population is *Nefm⁺/Mki67⁺/Prph⁺* and lacks *Chga/Chgb* expression, indicating a proliferative neuroblast-like cell population distinct from the surrounding *CHGA*-positive chromaffin PCC-tumor. **e** Unclassified *NF1*-cKO tumor (same case as Fig. 1h-i). H&E shows a small-cell neoplasm lacking classic PCC architecture. IF analyses (NEFM, CHGB, PRPH) demonstrate mixed chromaffin and neuroblast-associated features consistent with a hybrid sympathoadrenal tumor.

**Extended Data 6.**
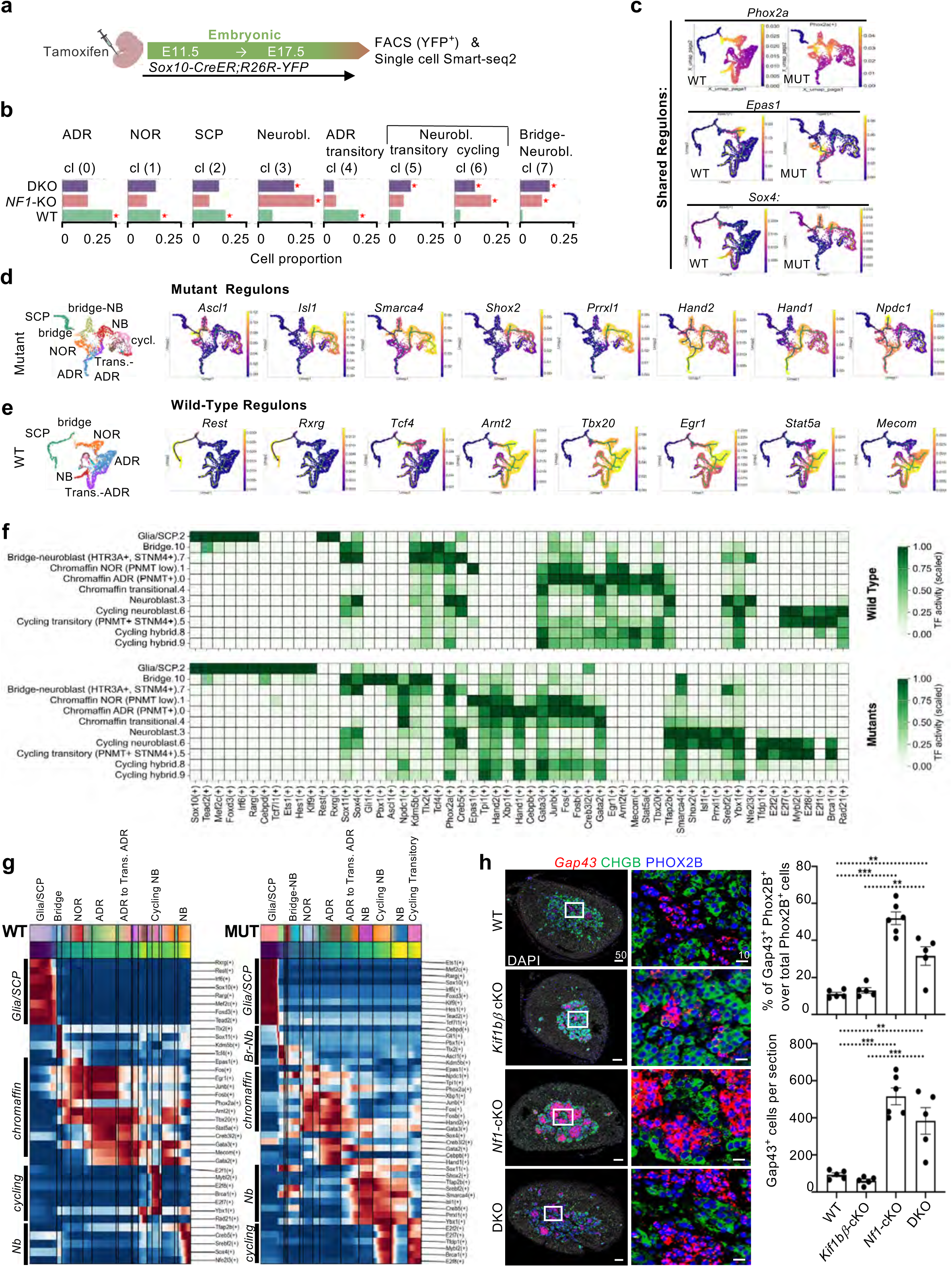
Extended overview of *NF1/KIF1Bb* loss inducing neurogenic regulatory programs along altered sympathoadrenal developmental trajectories. **a** Experimental design. Sox10-CreER;R26R-YFP embryos were pulse-labeled with tamoxifen at E11.5 and adrenal medullae harvested at E17.5. YFP⁺ cells were isolated by FACS and subjected to Smart-seq2 single-cell RNA sequencing. **b** Genotype-dependent shifts in cell state composition. Bar plots showing the percentage of cells per genotype within each transcriptional cluster. NF1-KO and DKO embryos exhibit reduced chromaffin representation and expansion of neuroblast and bridge-neuroblast populations compared to WT. **c-e** Feature plots of selective representative regulons mapped onto trajectory embeddings shown in main Fig. 2f-g. **c** Shared regulons active in both WT and mutant embryos include *Phox2a*, *Epas1* and *Sox4*. **d** Mutant-specific regulons include *Ascl1, Isl1, Smarca4, Shox2, Prrxl1, Hand1, Hand2,* and *Npdc1*, which are enriched along neuroblast and bridge-neuroblast trajectories and absent in WT embryos. **e** Wild-type specific regulons mapped onto wildtype trajectory that were associated with chromaffin differentiation and stabilization. **f** Global regulon activity across adrenal medullary cell states. Heatmap showing SCENIC regulon activity (AUCell scores) across E17.5 adrenal medullary cell states in wild-type (WT) and DKO mutant (Mut) embryos. Conserved lineage-associated regulons include Sox10, Tead2, and Foxd3 in SCP/glial cells; E2f family members and Mybl2 in proliferative states; Sox4 and Sox11 in neuroblast and bridge–neuroblast populations; and Epas1 in noradrenergic chromaffin cells. Mutant embryos show selective activation of additional neurogenic and chromatin-associated regulons. **g** Genotype-specific regulatory programs along developmental trajectories. Heatmaps showing transcription factor regulon activity mapped along inferred developmental trajectories in WT (left) and mutant (right) embryos, highlighting genotype-specific differences in regulatory state transitions. **h** RNAscope and immunofluorescence validation of neuroblast expansion. Representative combined RNAscope and immunofluorescence staining for *Gap43* (neuroblast marker), CHGB (chromaffin marker), and PHOX2B (sympathoadrenal lineage) with DAPI nuclear counterstain in E17.5 adrenal glands from WT, *KIF1Bβ*-cKO, *NF1*-cKO, and *NF1/KIF1Bβ*-DKO embryos. WT and *KIF1Bβ*-cKO adrenals are dominated by CHGB⁺ chromaffin cells with sparse *Gap43*⁺ neuroblasts. In contrast, *NF1*-cKO and DKO embryos display large multicellular clusters of *Gap43*⁺ neuroblasts. Insets show higher magnification views of boxed regions. Right: Quantification of neuroblast abundance measured as the proportion of *Gap43*⁺ cells or *Gap43*⁺/PHOX2B⁺ double-positive cells relative to total PHOX2B⁺ medullary cells as indicated. Both *NF1*-cKO and DKO embryos exhibit a significant expansion of neuroblasts states compared to WT and *KIF1Bβ*-KO controls.

**Extended Data 7.**
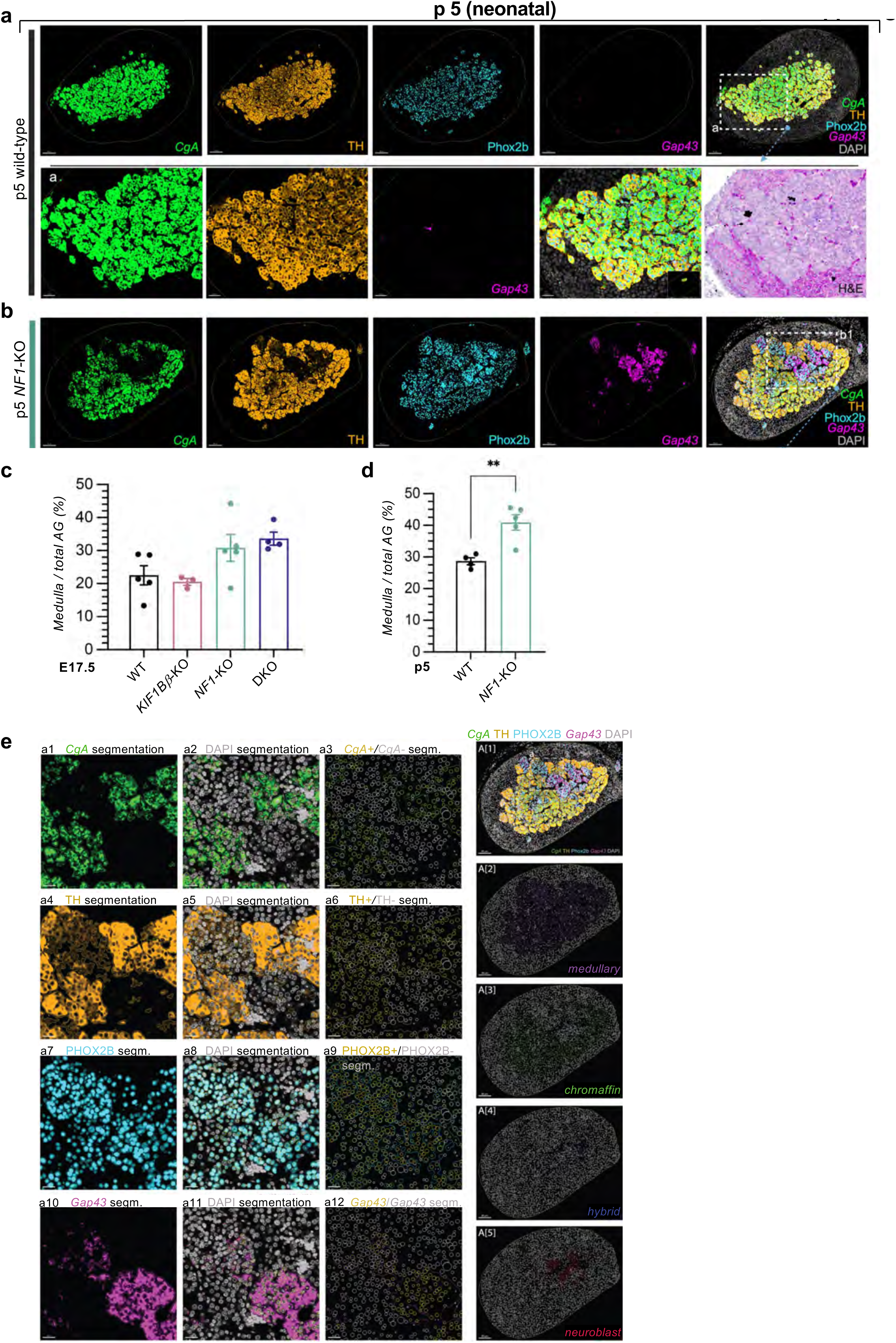
Persistent neonatal neuroblastic islands in *Nf1*-deficient medulla and segmentation-based classification of medullary cell populations. **a** Chromaffin-dominated adrenal medulla in wild-type neonates. Combined RNAscope *in situ* hybridization (*Chga, Gap43*) and immunofluorescence (PHOX2B, TH) on paraffin sections of postnatal day 5 (P5) adrenal glands from wild-type (WT) mice. Chromaffin cells are identified as PHOX2B⁺/*Chga⁺*/TH⁺/*Gap43^-^*, whereas neuroblasts are defined as PHOX2B⁺/Gap43⁺/*Chga^-^* with low or absent TH expression. WT adrenal medullae consist almost exclusively of mature chromaffin cells with only rare cells expressing neuroblast-associated markers. Insets (a) show higher magnification of the boxed region (scales, main and inset: 80μm and 30μm). Corresponding H&E staining of the same section is shown for histological reference. **b** Persistence of neuroblastic islands in Nf1-deficient adrenal medulla. Representative RNAscope/ immunofluorescence images of P5 adrenal glands from *Nf1*-KO mice reveal clusters of *Gap43⁺* neuroblasts embedded within the medullary compartment. Insets (b₁) show higher magnification views of the respective boxed regions shown in Figure 3a. **c-d** Expansion of the adrenal medulla in Nf1-deficient mice. Quantification of medullary area relative to the total adrenal gland area at embryonic day E17.5 (c) and postnatal day P5 (d). Nf1-KO adrenals display an enlarged medullary compartment compared with WT controls, indicating persistent expansion of the sympathoadrenal lineage after birth. Data represent mean ± SEM; each point represents one adrenal gland. **e** Representative workflow for segmentation and classification of chromaffin, neuroblast, and hybrid cell populations in adrenal gland sections. Panels a1–a12 illustrate sequential steps of signal detection, DAPI-based nuclear segmentation, and marker classification for *CgA*, TH, PHOX2B, and *Gap43* channels. Panels A1–A5 show whole-section maps illustrating the spatial distribution of medullary (PHOX2B⁺), chromaffin (*CgA⁺*/TH⁺), hybrid (*CgA⁺/Gap43⁺*), and neuroblast (*Gap43⁺*) cell populations across the adrenal gland. Cell identities were assigned using an Imaris-based pipeline based on marker co-expression and spatial proximity to segmented signal surfaces.

**Extended Data 8.**
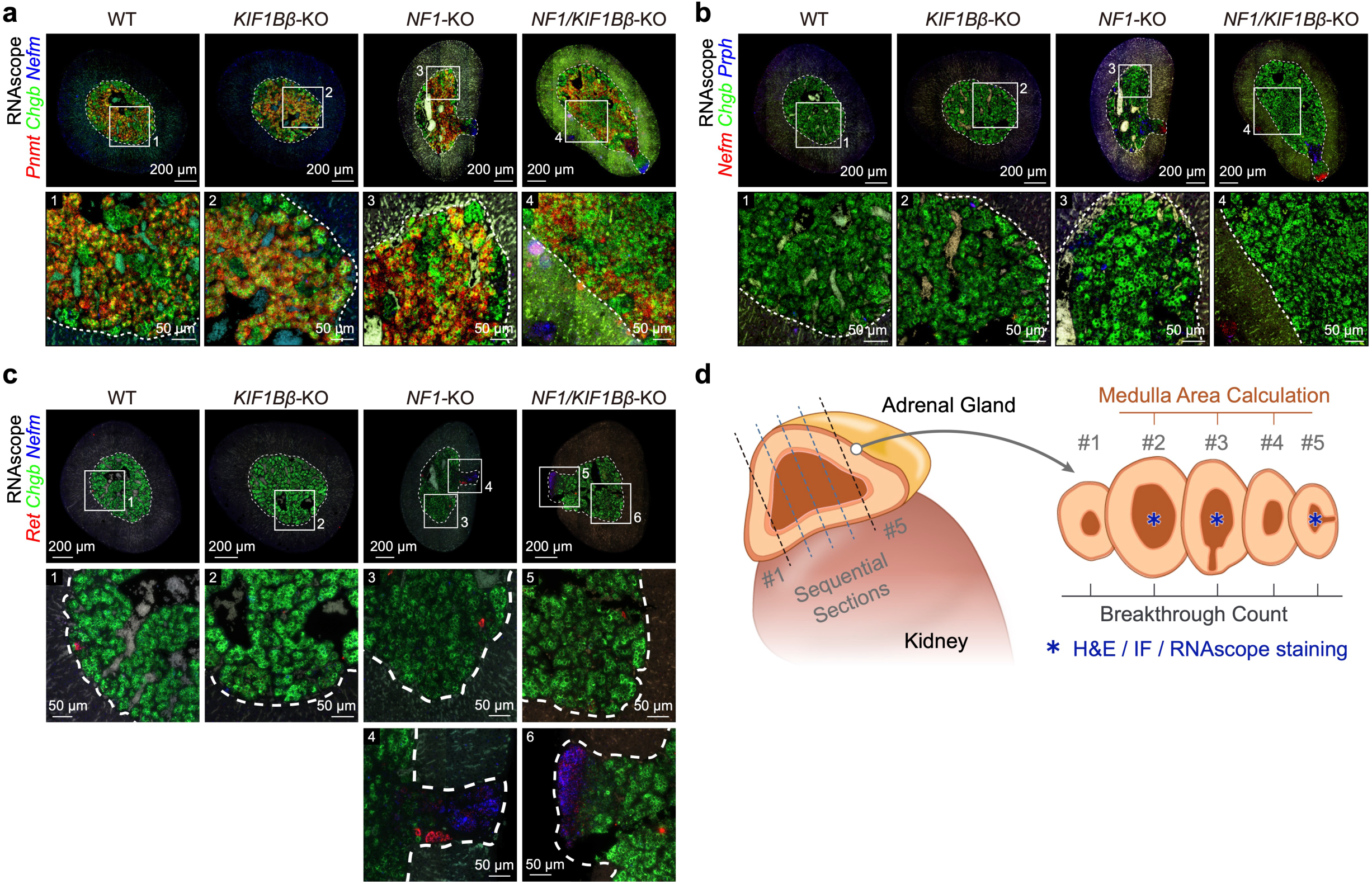
Preserved chromaffin identity in the adult medullary core and focal *Ret* upregulation in cortical breakthrough regions of mutant adrenals. **a-b** Chromaffin marker expression in the adult medullary core. RNAscope analysis of chromaffin markers (*Pnmt, Chgb*) together with neuroblast-associated genes (*Nefm, Prph*) in adrenal sections from 3-month-old wild-type (WT), *Kif1bβ*-KO, *Nf1*-KO, and *Nf1/Kif1bβ*-DKO mice. Across all genotypes, the medullary core retains strong chromaffin marker expression with only minimal and sporadic neuroblast marker expression, indicating preserved chromaffin differentiation within the central medullary compartment despite medullary hyperplasia. Neuroblast-associated programs are therefore not broadly reactivated within the adult medullary core and instead become enriched within focal cortical breakthrough regions described in Fig. 3k-n. Sections shown are adjacent serial sections to those presented in Fig. 3h-j. Scale bars: 200 µm (overview panels) and 50 µm (magnified views). **c** RNAscope analysis of *Ret*, *Chgb* and *Nefm* in adrenal sections from 3-month-old WT, *Kif1bβ*-KO, *Nf1*-KO and *Nf1/Kif1bβ*-DKO mice. *Ret* expression was low in the medullary core but increased within cortical breakthrough regions of *Nf1*-KO and DKO adrenals. Increased *Ret* expression was observed in cortical breakthroughs from three independent *Nf1*-KO and four independent DKO mice, coinciding spatially with regions in which chromaffin-to-neuroblast transitions were detected (Fig. 5). Scale bars: 200 µm (overview panels) and 50 µm (magnified views). **d** Serial sectioning strategy for histological and molecular analyses. Schematic illustrating the sequential sectioning approach used for histological (H&E), immunofluorescence, and RNAscope analyses and for quantification of medullary area and cortical breakthrough frequency. Consecutive sections (#1–#5) were used to maintain spatial alignment across staining modalities.

**Extended Data 9.**
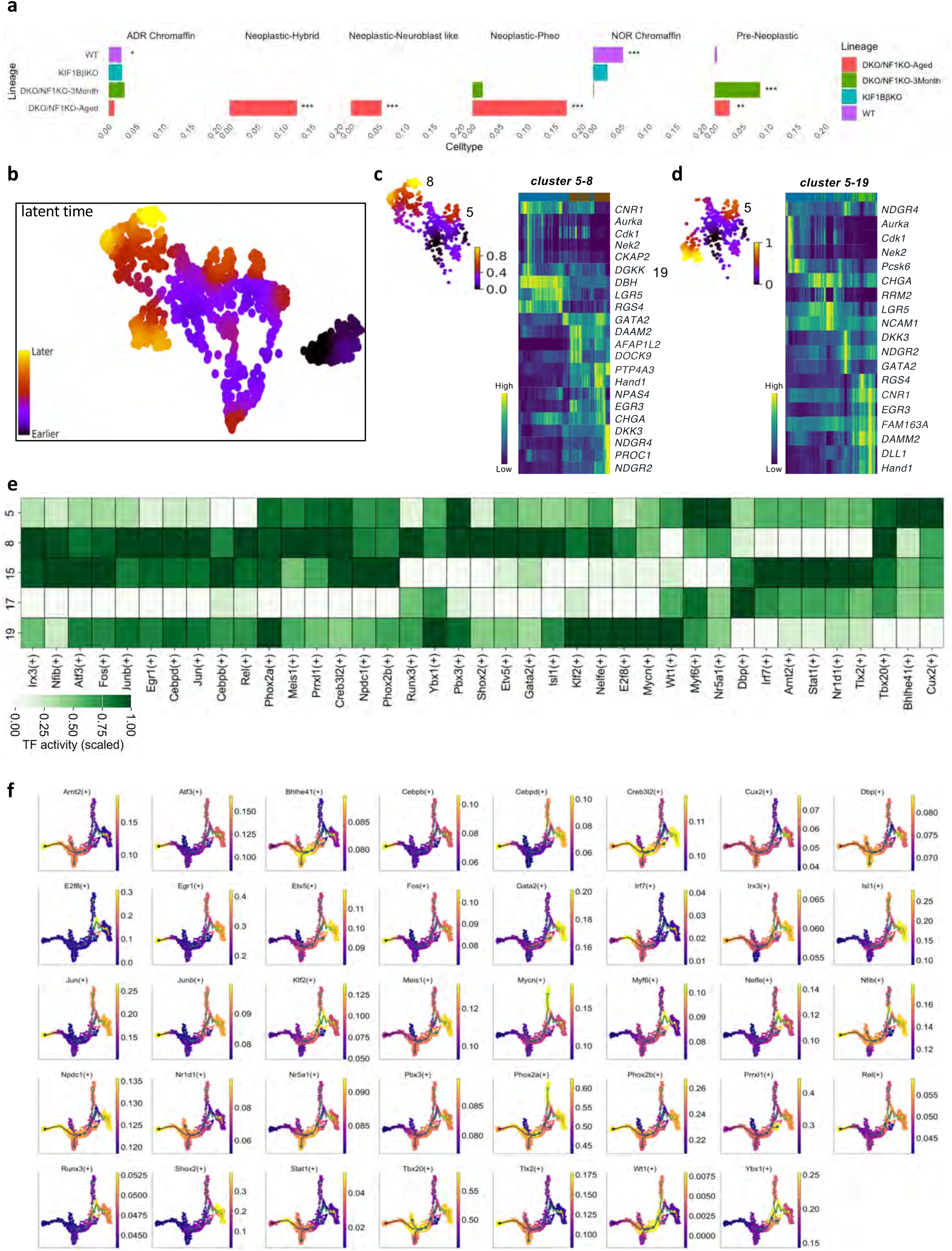
Extended analysis of cluster composition, transcriptional dynamics, and regulon activity associated with Fig. 6. **a** Cluster composition across genotypes and disease stages. Cluster proportion analysis across genotypes and disease stages corresponding to Fig. 5e. **b** Latent time progression across medullary tumor states. Latent time projection of medullary cells derived from RNA velocity analysis, illustrating inferred transcriptional progression across chromaffin, pre-neoplastic, and neoplastic cell states. **c-d** Gene expression dynamics along inferred tumor trajectories. Heatmaps showing genes differentially expressed along lineage transitions between tumor states, including transitions between hybrid and neuroblast-like populations (c) and between pheochromocytoma-like and neuroblast-like states (d). These analyses highlight progressive changes in neuroendocrine, neuronal, and proliferation-associated gene programs. **e** Regulon activity across medullary tumor clusters. Heatmap of SCENIC regulon activity (AUCell scores) across medullary clusters, showing chromaffin-associated regulatory programs together with tumor-state–specific activation of neurogenic and proliferation-associated transcription factors. **f** Regulon activity mapped along the inferred trajectory. Selected transcription factor regulons projected onto the trajectory structure shown in Fig. 5g-h. Neurogenic regulons, including *SHOX2* and *ISL1*, are enriched in hybrid tumor states, whereas a *MYCN*-centered regulatory program is selectively activated in neuroblast-like tumor cells. Core sympathoadrenal lineage regulons, including *PHOX2A* and *PHOX2B*, remain broadly active across tumor states.

**Extended Data 10.**
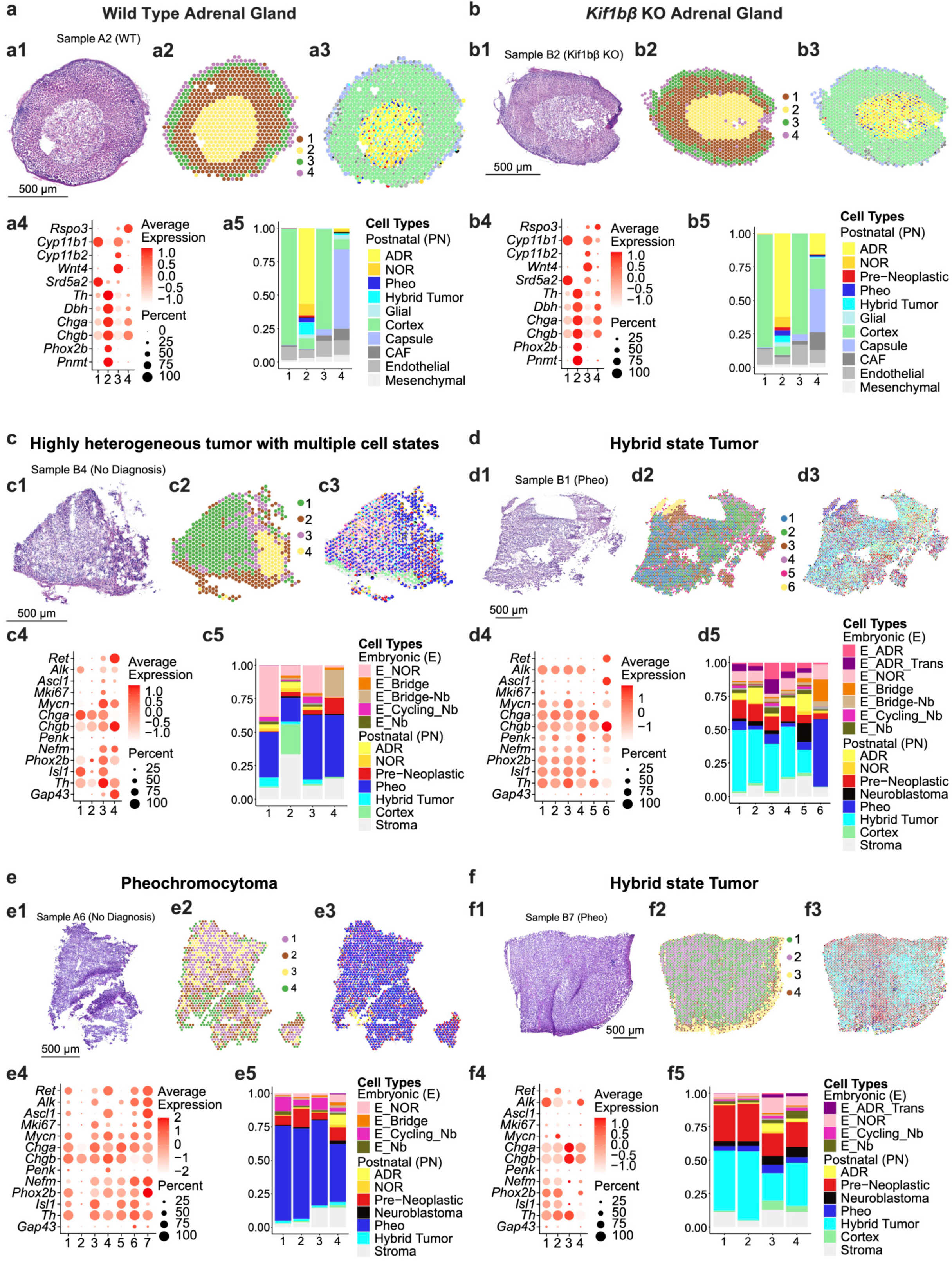
Spatial ISS recapitulates adrenal tissue architecture in non-neoplastic glands and reveals diverse tumor cell-state architectures. **a-b** Validation of spatial deconvolution in healthy adrenal glands. Targeted ISS was performed on adrenal glands from wild-type (WT) (a) and *Kif1bβ-*KO (b) mice using the 100-gene panel described in Fig. 6. Panels 1–5 follow the same layout as in Fig. 6. Panel 1 shows hematoxylin and eosin (H&E) staining for anatomical orientation. Panel 2 displays hexagonal spatial binning (∼40 μm “mini-bulks”) with Leiden clustering of transcriptomic profiles. Panel 3 shows spatial deconvolution using postnatal adrenal cell states as references. Panel 4 summarizes expression of selected marker genes (*Rspo3, Cyp11b1, Cyp11b2, Wnt4, Srd5a2, Th, Dbh, Chga, Chgb, Phox2b, Pnmt*) across spatial clusters. Panel 5 shows the proportional contributions of reference cell states within each cluster. Together, these analyses recapitulate the canonical capsule–cortex–medulla architecture, confirming accurate assignment of capsule, cortical, and chromaffin cell identities. **c-f** Additional tumor architectures revealed by spatial ISS. Tumor sections were analyzed using the same spatial workflow described in Fig. 6. Marker genes shown in panel 4 for tumor samples include *Ret, Alk, Ascl1, Mki67, Mycn, Chga, Chgb, Penk, Nefm, Phox2b, Isl1, Th*, and *Gap43*. **c** Highly heterogeneous tumor (sample B4, *Nf1/Kif1bβ*-DKO). Spatial deconvolution reveals a gradual transition from chromaffin-like regions to pheochromocytoma-like tumor states across the tissue. **d** Hybrid-dominant tumor (sample B1, *Nf1*-KO). Tumor regions are enriched for hybrid and pre-neoplastic states, with widespread expression of neurodevelopmental markers. **e** Uniform pheochromocytoma tumor (sample A6, *Nf1/Kif1bβ*-DKO). Spatial mapping shows consistent pheochromocytoma-like identity across the tumor. **f** Hybrid-dominant tumor (sample B7, *Nf1/Kif1bβ*-DKO). Spatial deconvolution reveals substantial hybrid and pre-neoplastic identities across the tumor, with smaller contributions from pheochromocytoma-, neuroblastoma- and embryonic developmental states.

## Supplementary Table legends

**Supplementary Table 1. Tumor penetrance and latency in *Kif1bβ/Nf1* double knockout (DKO) and *Nf1* conditional knockout mice.**

Penetrance is reported as the percentage of tumor-bearing animals. Latency and median survival are given in days. Censored animals were excluded from survival analysis.

**Supplementary Table 2: Summary of histopathological tumor diagnoses in *Kif1bβ/Nf1* double knockout (DKO) and *Nf1* conditional knockout mice.**

Tumor diagnoses were assigned based on H&E histopathological evaluation and reviewed by expert endocrine pathologists. Tumors were categorized as pheochromocytoma (intra-adrenal), pheochromocytoma with focal small blue cell component, composite pheochromocytoma–neuroblastoma, paraganglioma (extra-adrenal), neuroblastoma, or unclassified tumor. Bilateral pheochromocytomas were counted as two independent tumors.

## Supplementary Data legends

**Supplementary Data 1**: **Human sample metadata and single-nucleus RNA sequencing summary.** Clinical information and single-nucleus RNA sequencing statistics for all human samples included in this study, comprising neuroblastoma, pheochromocytoma and paraganglioma (PPGL), and normal adrenal gland samples. Neuroblastoma samples derived from the same patient are listed consecutively and separated from those of different patients by a blank row. NED, no evidence of disease; AWD, alive with disease; DOD, died of disease; WT, wild type; MUT, somatic mutation; MUT (const), constitutional (germline) mutation.

**Supplementary Data 2**: **Differentially expressed genes for each cell type identified in the merged NB and PPGL single-nucleus RNA sequencing dataset.** Each worksheet contains the differentially expressed genes for one cell type. Differential expression analysis was performed using the Wilcoxon rank-sum test. P values were adjusted for multiple testing using the Bonferroni correction.

**Supplementary Data 3: Differentially expressed genes for each cell type identified in the human PPGL #1 spatial transcriptomics dataset.** Each worksheet contains the differentially expressed genes for one cell type. Differential expression analysis was performed using the Wilcoxon rank-sum test. P values were adjusted for multiple testing using the Bonferroni correction.

**Supplementary Data 4: Differentially expressed genes for each cell state identified in the human PPGL #1 spatial transcriptomics dataset.** Each worksheet contains the differentially expressed genes for one cell type. Differential expression analysis was performed using the Wilcoxon rank-sum test. P values were adjusted for multiple testing using the Bonferroni correction.

**Supplementary Data 5. Shared differentially expressed genes defining corresponding cell states in the fetal adrenal reference and human PPGL #1 spatial transcriptomics dataset.** Each worksheet contains the shared differentially expressed genes for one corresponding cell state. Differential expression statistics from the fetal adrenal reference and the composite human pheochromocytoma spatial transcriptomics datasets are reported separately, with column names suffixed by “_Ref” and “_Tumor”, respectively. P values were calculated using the Wilcoxon rank-sum test and adjusted for multiple testing using the Bonferroni correction. pct.1 and pct.2 indicate the proportions of cells expressing the indicated gene in the target cell state and in all remaining cell states, respectively.

**Supplementary Data 6. Differentially expressed genes for each cell type identified in the malignant PPGL #263 spatial transcriptomics dataset.** Each worksheet contains the differentially expressed genes for one cell type. Differential expression analysis was performed using the Wilcoxon rank-sum test. P values were adjusted for multiple testing using the Bonferroni correction.

**Supplementary Data 7. Differentially expressed genes for each cell state identified in the malignant PPGL #263 spatial transcriptomics dataset.** Each worksheet contains the differentially expressed genes for one cell type. Differential expression analysis was performed using the Wilcoxon rank-sum test. P values were adjusted for multiple testing using the Bonferroni correction.

**Supplementary Data 8. Sample metadata and sequencing statistics for postnatal and embryonic (E17) adrenal single-cell RNA-seq datasets.** Overview of mouse adrenal gland samples across genotypes and developmental stages. For each sample, the table reports library identifiers, genotype/stage, number of cells sequenced, total number of reads, number of high-quality (HQ) cells retained after filtering, and number of reads used for downstream cell clustering.

**Supplementary Data 9. Differentially expressed genes for each cell type in the E17 mouse sympathoadrenal dataset.** Each worksheet contains genes identified by comparing the indicated cell type with all remaining cell types using the Wilcoxon rank-sum test. P values were calculated using the Wilcoxon rank-sum test and adjusted for multiple testing using the Bonferroni correction.

**Supplementary Data 10. Differentially expressed genes for each cell type in the postnatal mouse adrenal gland and tumor dataset.** Each worksheet contains genes identified by comparing the indicated cell type with all remaining cell types using the Wilcoxon rank-sum test. P values were calculated using the Wilcoxon rank-sum test and adjusted for multiple testing using the Bonferroni correction.

**Supplementary Data 11. Selected Gene Panel for In-Situ Sequencing (ISS).** Gene panel for in situ sequencing comprising 100 genes selected from markers and differentially expressed genes across embryonic, postnatal, pre-neoplastic, and tumor cell clusters identified in single-cell RNA-seq datasets (Supplementary Data 9 and 10).

**Supplementary Data 12. Cluster-specific marker genes of pseudo-spot clusters identified from mouse adrenal gland and tumor in situ sequencing (ISS) data.** Marker genes were identified using a Gini coefficient–based approach implemented in Giotto (findMarkers_one_vs_all), comparing each cluster to all other clusters. Genes are ranked based on combined expression and detection specificity, as defined by Gini scores. Only genes with a minimum expression Gini score ≥ 0.1 and detection Gini score ≥ 0.1 were retained. This method does not involve statistical hypothesis testing or multiple testing correction. Reported markers therefore represent genes with high cluster-specific expression and detection specificity.

