## Supplementary Table 1-2 and legends of Supplementary Data 1-12 for "Spatially organized developmental cell states link paraganglioma and neuroblastoma through chromaffin-neuroblast plasticity"

**Supplementary Information: Document S1:**

**Page 2-3: Supplementary Tables 1-2 including legends**

**Page 4-7: Legends of Supplementary Data 1-12**

| Genotypes | KIF1Bb NF1 DBHCre | NF1 DBHCre |
|---|---|---|
| Number of Animals | 11 | 20 |
| Penetrance(%) | 100 | 100 |
| Latency and Median survival (days) | 140, 407 | 224, 496 |

**Supplementary Table 1. Tumor penetrance and latency in *Kif1bβ/Nf1* double knockout (DKO) and *Nf1* conditional knockout mice.**

Penetrance is reported as the percentage of tumor-bearing animals. Latency and median survival are given in days. Censored animals were excluded from survival analysis.

| Genotypes | KIF1Bb NF1 ko | NF1 ko |
|---|---|---|
| Number of Animals | 11 | 20 |
| Number of tumors | 21 | 42 |
| Pheochromocytoma | 14 | 29 |
| Pheochromocytoma with small, blue cells | 4 | 6 |
| Composite Pheochromocytoma with Neuroblastoma | 2 | 1 |
| Unclasified tumor with small, blue cells | 0 | 1 |
| Paraganglioma | 0 | 5 |
| Neuroblastoma | 1 | 0 |

**Supplementary Table 2: Summary of histopathological tumor diagnoses in *Kif1bβ/Nf1* double knockout (DKO) and *Nf1* conditional knockout mice.**

Tumor diagnoses were assigned based on H&E histopathological evaluation and reviewed by expert endocrine pathologists. Tumors were categorized as pheochromocytoma (intra-adrenal), pheochromocytoma with focal small blue cell component, composite pheochromocytoma–neuroblastoma, paraganglioma (extra-adrenal), neuroblastoma, or unclassified tumor. Bilateral pheochromocytomas were counted as two independent tumors.

**Supplementary Data legends:**

**Supplementary Data 1. Human sample metadata and single-nucleus RNA-sequencing summary.** Clinical information and single-nucleus RNA sequencing statistics for all human samples included in this study, comprising neuroblastoma, pheochromocytoma and paraganglioma (PPGL), and normal adrenal gland samples. Neuroblastoma samples derived from the same patient are listed consecutively and separated from those of different patients by a blank row. NED, no evidence of disease; AWD, alive with disease; DOD, died of disease; WT, wild type; MUT, somatic mutation; MUT (const), constitutional (germline) mutation.

**Supplementary Data 2. Differentially expressed genes for each annotated cell type in single-nucleus RNA-sequencing analyses of human NB and PPGL.** Each worksheet contains genes identified by comparing the indicated cell type with all remaining cell types using the Wilcoxon rank-sum test. *P* values were adjusted for multiple testing using the Bonferroni correction.

**Supplementary Data 8. Sample metadata and sequencing statistics for postnatal and embryonic (E17) mouse adrenal single-cell RNA-sequencing datasets.** Overview of mouse adrenal gland samples across genotypes and developmental stages. For each sample, the table reports the library identifier, genotype or developmental stage, number of cells sequenced, total number of reads, number of high-quality (HQ) cells retained after filtering, and number of reads retained for downstream analysis.

**Supplementary Data 9. Differentially expressed genes for each cell type in the E17 mouse sympathoadrenal dataset.** Each worksheet contains genes identified by comparing the indicated cell type with all remaining cell types using the Wilcoxon rank-sum test. *P* values were adjusted for multiple testing using the Bonferroni correction.

**Supplementary Data 10. Differentially expressed genes for each cell type in the postnatal mouse adrenal gland and tumor dataset.** Each worksheet contains genes identified by comparing the indicated cell type with all remaining cell types using the Wilcoxon rank-sum test. *P* values were adjusted for multiple testing using the Bonferroni correction.

**Supplementary Data 11. Selected Gene Panel for In-Situ Sequencing (ISS).** List of 100 genes selected based on marker and differential-expression analyses of embryonic, postnatal, preneoplastic and tumor cell populations identified in the single-cell RNA-sequencing datasets (Supplementary Data 9 and 10).

**Supplementary Data 12. Cluster-specific marker genes of pseudo-spot clusters identified from mouse adrenal gland and tumor in situ sequencing (ISS) data.** Marker genes were identified using a Gini coefficient–based approach implemented in Giotto (findMarkers_one_vs_all), comparing each cluster to all other clusters. Genes are ranked based on combined expression and detection specificity, as defined by Gini scores. Only genes with a minimum expression Gini score ≥ 0.1 and detection Gini score ≥ 0.1 were retained. This method does not involve statistical hypothesis testing or multiple testing correction. Reported markers therefore represent genes with high cluster-specific expression and detection specificity.
